# The response of *Allolobophora chlorotica* to drought stress in four soils

**DOI:** 10.1101/2025.11.06.686976

**Authors:** R. V. A. Bray, M. E. Hodson, P. J. Watt

## Abstract

Earthworms are key contributors to healthy and productive soils, yet their reliance on water makes them vulnerable to the increased frequency and severity of droughts predicted under climate change. To avoid desiccation, some earthworms induce aestivation, a period of reduced metabolism during which they coil up and seal themselves into a chamber until conditions improve. However, the environmental conditions that trigger aestivation remain poorly understood. Here the responses of *Allolobophora chlorotica*, a common UK earthworm, to gradual air drying (at 15 ± 1 °C) were examined in four soils differing in texture. Earthworm activity (active or aestivating) and mass change relative to initial values (pre- and post-24 hours hydration) were measured at three gravimetric moisture contents (~19.7, 15.55 and 12.39 wt%) and three water potentials (~pF 1.59, 2.92 and 4.05). Water potential, rather than bulk water content, was the strongest predictor of behaviour. All earthworms remained active and gained ~36-53 % mass at the highest water availability (~pF 1.59), but 100 % aestivated at the lowest (~pF 4.05) in all but the sandiest soil. In contrast, responses at equal gravimetric moisture contents varied by soil type. All individuals in the clay aestivated and lost up to ~45 % mass, whereas those in sand and sandy loam soil largely remained active and gained mass. Differences likely reflect textural constraints on movement and the construction of aestivation chambers, which were fragile in sandy soils but more robust in clay-rich soils. After 24 hours of hydration, all earthworms had increased beyond their starting mass, indicating changes in mass were largely due to reversible water loss. However, some residual mass differences between control and drying treatments suggest differences in tissue mass, potentially attributable to suspended feeding and clitellum regression, characteristic features of aestivation. Overall, these findings show that *Al. chlorotica* is highly desiccation tolerant, but that soil texture strongly modulates both the onset and viability of aestivation, with implications for predicting earthworm resilience under future drought regimes.

## Introduction

One well-documented consequence of climate change is the increasing frequency and severity of droughts (Seneviratne *et al*., 2021). These events pose a major challenge for soil organisms such as earthworms, which depend on free water for critical functions such as facilitating oxygen uptake for respiration (Holmstrup, 2001), and forming coelomic fluid, which acts as a hydraulic skeleton to maintain turgor and permit movement through the soil (Kretzschmar and Bruchou, 1991; Ramsay, 1949; Whalen *et al*., 2000). Under water-limiting conditions, some earthworms enter aestivation, a period of suspended development (Jiménez *et al*., 2000) characterised by coiling to reduce surface exposure to the soil (McDaniel *et al*., 2013a), expulsion of gut contents (Holmstrup, 2001), and construction of a chamber in which they enclose themselves (Bayley *et al*., 2010). During aestivation, metabolic activity is reduced and their role in the provision of ecosystem services as key ecosystem engineers is halted until conditions improve (Gerard, 1967; Jiménez *et al*., 2000; McDaniel *et al*., 2013b).

Soil water is commonly described by gravimetric or volumetric moisture content, which quantify water per unit dry soil mass or volume, respectively (Weil and Brady, 2017). However, soil water potential is often considered to be a more biologically relevant measure, as it reflects the force at which water is held in the soil and therefore the energy required for organisms to extract it (Lee, 1985; Kretzschmar and Bruchou, 1991; McDaniel *et al*., 2013a). Water potential, expressed in hectopascals (hPa), kilopascals (kPa) or as a pF value where pF = log(hPa), increases as soils dry and water becomes less available. Field capacity typically occurs at pF 2 (10 kPa) and represents the maximum amount of water held by soil after excess has drained away (Weil and Brady, 2017). Above pF 3.5, free water is typically no longer readily available in soil (Friis *et al*., 2004), and at pF 4.2 (1,500 kPa), the permanent wilting point is reached, beyond which plants cannot recover their turgidity, a common occurrence in temperate regions during summer (Friis *et al*., 2004; Weil and Brady, 2017). The relationship between soil moisture content and water potential is strongly dependent on soil texture, meaning soils can differ substantially in their water availability despite having the same water content (Kretzschmar and Bruchou, 1991; Collis-George, 1959). Generally, at the same moisture content and mass of dry soil, finer-textured soils such as clays have a larger surface area and hold water more tightly than coarser soils like sands and so have a lower water availability (Potvin and Lilleskov, 2017; Weil and Brady, 2017).

Determining the soil water thresholds that trigger changes in earthworm behaviour is essential to predict how drought will affect their activity, survival, and the ecosystem services they support. Previous studies indicate that earthworms remain active above 20 wt% soil moisture (Díaz Cosín *et al*., 2006; Tilikj and Novo, 2022), but that their activity is constrained at water potentials above pF 3 to 3.3 (Gerard, 1967; Kretzschmar and Bruchou, 1991; Nordström, 1975; Rundgren, 1975; Baker *et al*., 1993). However, available data is relatively scarce, tolerances are likely to be species-specific, and further work is required to improve our understanding of the precise conditions that induce aestivation.

This study examined how soil moisture content and water potential influence the responses of *Allolobophora chlorotica*, one of the most common UK earthworms (Ashwood *et al*., 2024), to drought stress in four soils of differing texture. Earthworms were maintained in soils that air-dried gradually, and individuals were sampled at the same three gravimetric moisture contents and the same three water potential values, derived from water retention curves for all four soils. At each sampling point, earthworms were categorised as active or aestivating and changes in body mass were recorded to assess physiological responses to drought and identify when conditions became limiting (Kretzschmar and Bruchou, 1991). We predicted that (i) the incidence of aestivation and mass loss would increase as soil water loss increased, and (ii) aestivation and earthworm mass loss would occur at higher gravimetric water contents in clay-rich soils due to their lower water availability at a given moisture content. If water availability governs earthworm responses to drought, then similar behavioural and physiological thresholds should be observed across soils at equivalent water potentials.

## Methods

### Soil types and properties

Four standard soils (LUFA 2.1, 2.2, 2.4 and 6S) were obtained from LUFA Speyer (Speyer, Germany). These soils are routinely used in soil ecology studies and were collected in 2022 from agricultural fields that had been free from pesticide, biocidal fertiliser and organic manure applications for at least five years (LUFA, Speyer 2022). Soils were air-dried, sieved to 2 mm and subsamples were oven-dried at 105 °C to constant mass to determine their residual moisture contents. Soil organic matter (SOM) was estimated as loss on ignition by burning oven-dried soil at 400 °C for 4 hours in a muffle furnace (following Jensen *et al*., 2018, but with reduced temperature to minimise structural water loss). Soil pH was measured from a suspension of 10 g air dried soil in 25 ml deionised distilled water (DIW), shaken end-over-end at 30 rpm for 15 minutes (The Analysis of Agricultural Materials – Ministry of Agriculture, Fisheries and Food). Measurements were made using a SciQuip 930 Precision pH/Ion meter calibrated with standards of pH 10, 7 and 4. SOM and pH were measured both for soils with and without sheep manure added as a food source (**Table 1**). Additional details of the soil compositions and properties can be found in **Table 1** and **Fig. S1**.

**Table 1.**
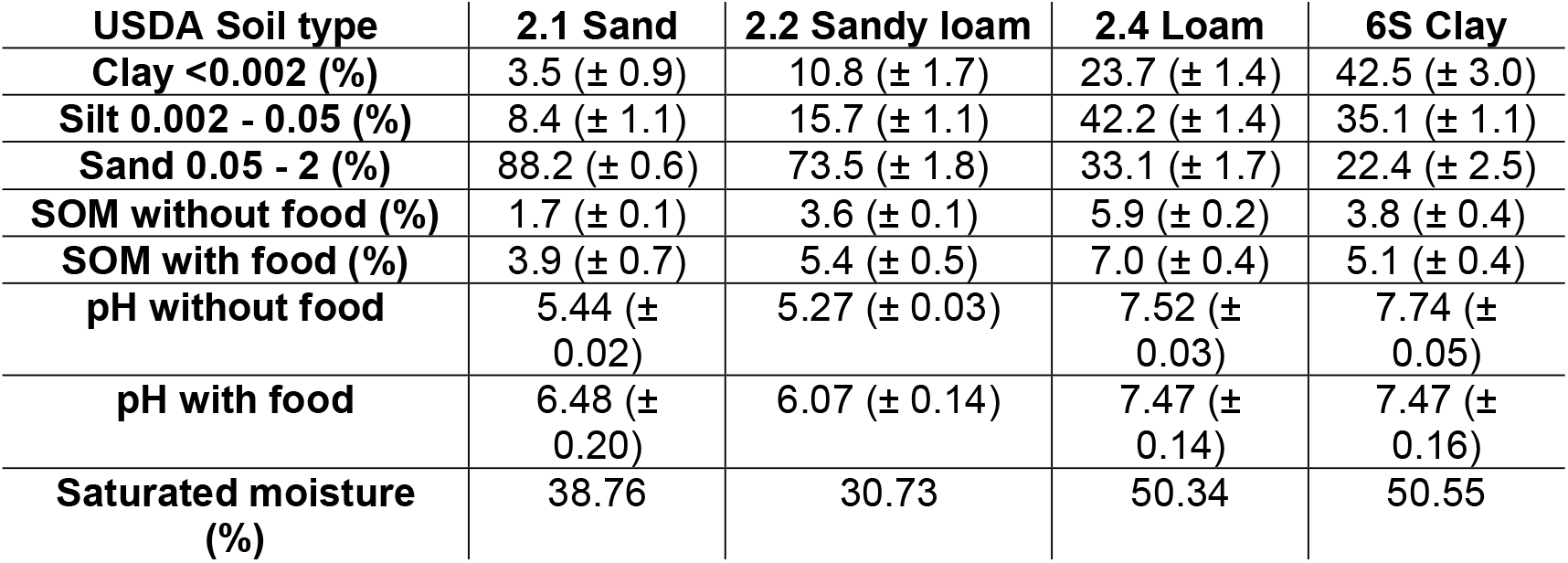
The particle size distribution (mm) % based on USDA data given in ‘Chemical and physical characteristics of standard soils’ (LUFA Speyer, 2022). Means with standard deviations given in brackets. The soil pH and SOM values are given both with (n = 50) and without (n = 3) sheep manure added as a food source. Saturated moisture content (vol%) taken from HYPROP measurements.

### Water retention curves

Soil water retention curves (**Fig. 1**) were generated using HYPROP 2 instruments (UMS 2015, **Fig. S2**). This system uses tensiometers to measure changes in the mass and water potential as saturated soil samples air-dry (see Supplementary Materials for procedural details). The HYPROP outputs volumetric soil moisture values, which were converted to gravimetric moisture contents using measured soil bulk densities to enable direct measurement of moisture content during the experiment.

**Figure 1.**
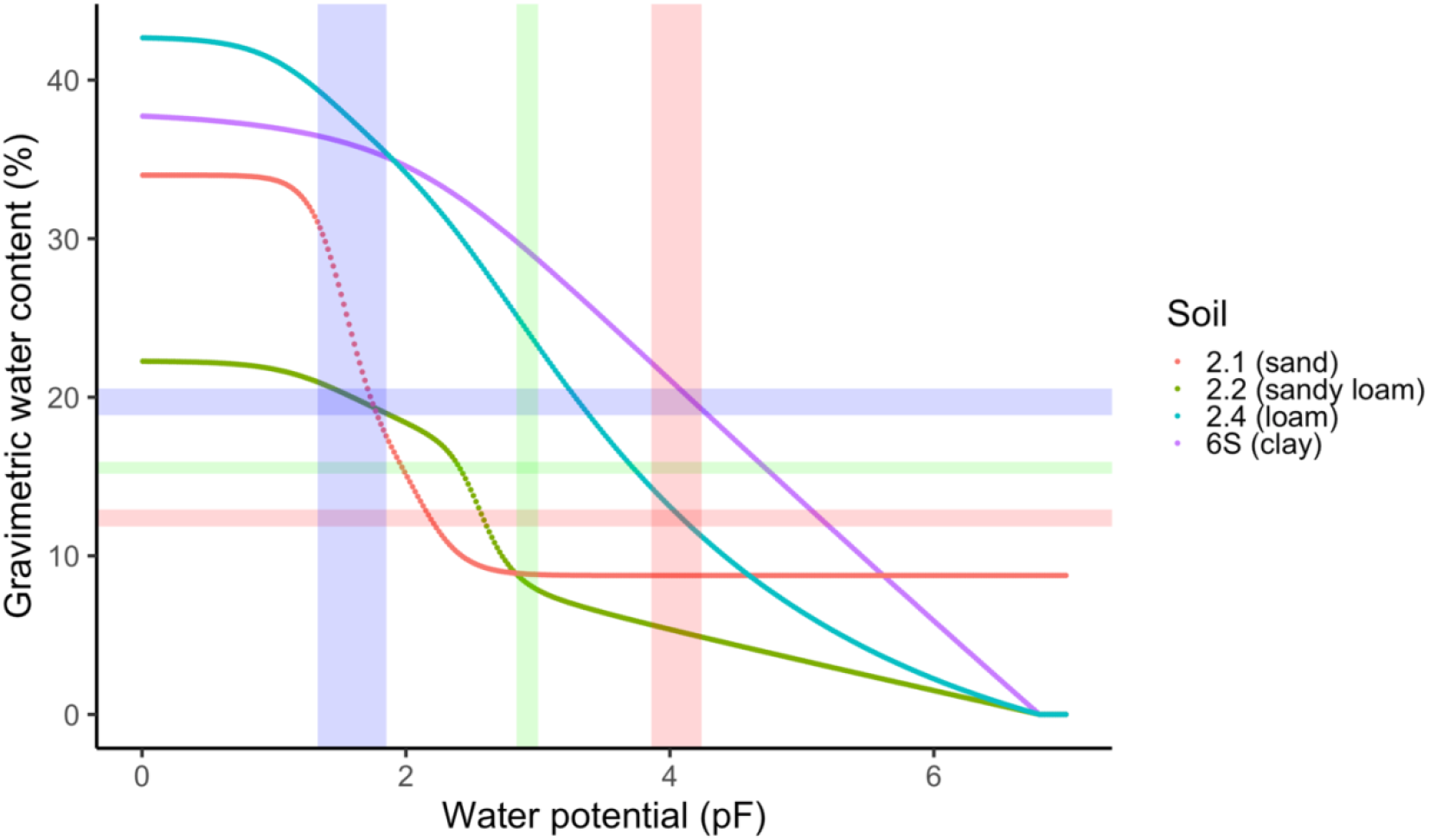
Four water retention curves showing the relationship between water potential (pF) and moisture content (wt%) for each of the standard soils 2.1, 2.2, 2.4 and 6S. Data is generated from the HYPROP values with curves fitted based on the PDI-variant of the binomial constrained van Genuchten model (m = 1 − 1/n). Shaded bands represent pF (vertical) and gravimetric water content (horizontal) sampling ranges (means ± standard error) determined post-sampling (blue = highest wt% and most available pF, green = intermediate sampling points, red = lowest wt% and least available pF).

### Earthworm collection and culturing

Adult (clitellate) *Allolobophora chlorotica* were collected in Spring 2024 by hand-sorting soil from pits (~18 cm x 18 cm at the surface to 20 cm depth) dug in the field margins of Warren Paddock, University of Leeds Research Farm, Tadcaster (53°52’25.9”N 1°19’33.4”W). The field had been in permanent pasture from 2012 and cut for silage until sown with oilseed rape in Autumn 2023. Earthworms were maintained in a controlled temperature room (15 ± 1 °C and in complete darkness) in field soil, a well-drained calcareous loam soil (Holden *et al*., 2019) with a pH of ~6.9 and SOM of ~8.2 % (determined in the laboratory following the same procedures noted previously). Soil moisture was maintained at ~25 wt% and earthworms were fed *ad-libitum* with sheep manure until experiments commenced.

### Experimental setup and sampling

Sheep manure (from sheep reared without the use of anti-helminthic drugs) was collected from a farm in Sheffield, oven-dried, ground to < 2 mm and added to the experimental soils as a food source. The fine particle size was chosen because smaller particle sizes of organic matter are easier for earthworms to ingest with Lowe and Butt (2003) finding *Al. chlorotica* attained a significantly higher mass (185 % greater) when provided with milled compared to unmilled separated cattle solids. Approximately 3.5 g of dry manure was incorporated into the soil (~0.5 g earthworm^-1^ week^-1^ as in Lowe and Butt, 2005) to provide enough food for 7 weeks. Soil and manure were homogenised and packed into 300 ml plastic cups, levelled to the 150 ml mark to achieve bulk densities consistent with the HYPROP cores: 2.1 = 1.14 g/cm^3^ (171 g dry mass), 2.2 = 1.38 g/cm^3^ (207 g dry mass), 2.4 = 1.18 g/cm^3^ (177 g dry mass), and 6S = 1.34 g/cm^3^ (201 g dry mass). Distilled deionised water (DIW) was applied using a spray bottle to reach initial gravimetric moisture contents of 22 % (2.1, 2.2), 38 % (2.4) and 37 % (6S), corresponding to water potentials between pF 0.8-1.6. These values were above the first sampling point but below saturation, maintaining high water availability while preserving soil structure. Control soils were kept at these moisture levels throughout.

*Al. chlorotica* (n = 200) were depurated on filter paper moistened with DIW for 24 hours, blotted dry, and weighed to determine fresh masses. Individuals were randomly allocated to drying (n = 100) or control (n = 100) treatments, with 25 earthworms per soil type. Mean initial earthworm fresh mass was 0.291 g (± 0.074 g), with no significant differences between treatments (ANOVA: F = 2.448, df = 1, 192, p = 0.119) or soil types (ANOVA: F = 1.066, df = 3, 192, p = 0.365) (**Fig. S3**). One earthworm was placed on the soil surface of a cup and the total mass was recorded to allow monitoring of water loss and addition of DIW to maintain water content in controls when needed. Cups were covered with nylon mesh, secured with an elastic band to allow evaporation but prevent earthworm escape. All cups were kept in complete darkness in a controlled temperature room at 15 ± 1 °C. Soils were maintained at constant moisture for 48 hours to allow acclimation, after which drying treatments received no additional water.

Drying treatments were sampled at three gravimetric moisture contents (~19.7, 15.55 and ~12.39 wt%) and at three water potentials (pF ~1.59, ~2.92 and ~4.05) (**Fig. 1, Table 2**). These gravimetric moisture contents were selected based on a prior range-finding test showing *Al. chlorotica* remained active down to 18 wt%, but most had entered aestivation by 13 wt% (unpublished results). The three water potential values were chosen to be close to field capacity, permanent wilting point and an intermediate water availability between the two. As each soil had a distinct water retention curve, identical gravimetric contents represented different water potentials and vice versa. In each soil type one of the pF levels corresponded to one of the gravimetric sampling points (**Fig. 1**) so that the total number of sampling points per soil was five rather than six.

**Table 2.** Mean and standard error value (n =20) for each of the a) gravimetric and b) water potential values determined after sampling.

| a) Gravimetric sampling points | Relative water content | b) Water potential sampling points | Relative water availability |
| --- | --- | --- | --- |
| 19.7 (± 0.84) wt% | Highest | pF 1.59 (± 0.26) | Most |
| 15.55 (± 0.38) wt% | Intermediate | pF 2.92 (± 0.08) | Intermediate |
| 12.39 (± 0.53) wt% | Lowest | pF 4.05 (± 0.19) | Least |

At each sampling point, five replicates per soil were destructively sampled from both the drying and control treatments. Soils were hand-sorted and *Al. chlorotica* were classified as active (extended, moving) or aestivating (coiled within a soil chamber). After gently removing excess soil, earthworms were weighed to obtain their pre-hydration mass, then placed on moist filter paper for 24 hours, blotted, and re-weighed to obtain their post-hydration mass.

### Statistical analyses

Analyses were conducted in R v4.4.1 (R Core Team, 2024). Diagnostic plots confirmed linearity, constant variance, and normality assumptions. Initial earthworm masses were not normally distributed and so were log transformed prior to ANOVA testing. Binomial glms (logit link) with main and interactive effects of soil type and soil water were fitted separately for the gravimetric and the water potential sampling points to look for differences in the proportion of aestivating *Al. chlorotica*. Changes in earthworm mass were calculated as percentage change relative to their initial mass. Log-transformed earthworm masses were analysed using ANOVA with interacting effects of soil type, state (active or aestivating), and measurement time (pre- and post-hydration) to look for significant differences in mass between those found active and aestivating. Two-way ANOVAs assessed effects of soil type and water (wt% or pF) on mass changes within drying treatments (both pre- and post-hydration). Three-way ANOVAs included treatment (drying vs. control) as an additional factor for between-group comparisons. Continuous relationships between earthworm mass change (pre- and post-hydration) and soil water availability (pF) or moisture (wt%) were explored using generalised additive models (GAMs) to examine non-linear relationships. For analyses involving ANOVAs, post-hoc treatment contrasts were performed using Tukey tests to identify which groups differed significantly. The rate of drying (water loss/day) was similar across soils (0.9-1 g day^-1^) and so was not included as a covariate in the models.

## Results

### State of Allolobophora chlorotica on sampling

Across all soil types, 99 % of *Al. chlorotica* in the constant-moisture control conditions remained active throughout the experiment. In contrast, under drying conditions, there was a significant effect of water potential on the proportion of aestivating individuals (Binomial distribution glm: X±^2^ = 30.468, df = 2, 9, p < 0.001), which varied significantly with soil type (Binomial distribution glm: X^2^ = 21.299, df = 3, 6, p < 0.001). All earthworms were active at the highest water availability (~pF 1.59). At the intermediate availability (~pF 2.92), aestivation occurred only in the sandier soils (2.1 and 2.2), while in the loam and clay soils (2.4 and 6S), aestivation was only observed at the lowest water availability (~pF 4.05) (**Fig. 2**). When data were expressed in terms of gravimetric moisture content, the proportion of aestivating individuals was marginally influenced by soil moisture (Binomial distribution glm: X^2^ = 5.858, df = 2, 9, p = 0.053) but again varied significantly with soil type (Binomial distribution glm: X^2^ = 50.332, df = 3, 6, p < 0.001). No individuals aestivated in the sand and sandy loam soils (2.1 and 2.2) until the lowest moisture level (~12.39 wt%), whereas aestivation was observed in the loam and clay soils (2.4 and 6S) even at the highest moisture content (~19.7 wt%) (**Fig. 3**). All earthworms in the clay soil were aestivating at each of the three gravimetric sampling points.

**Figure 2.**
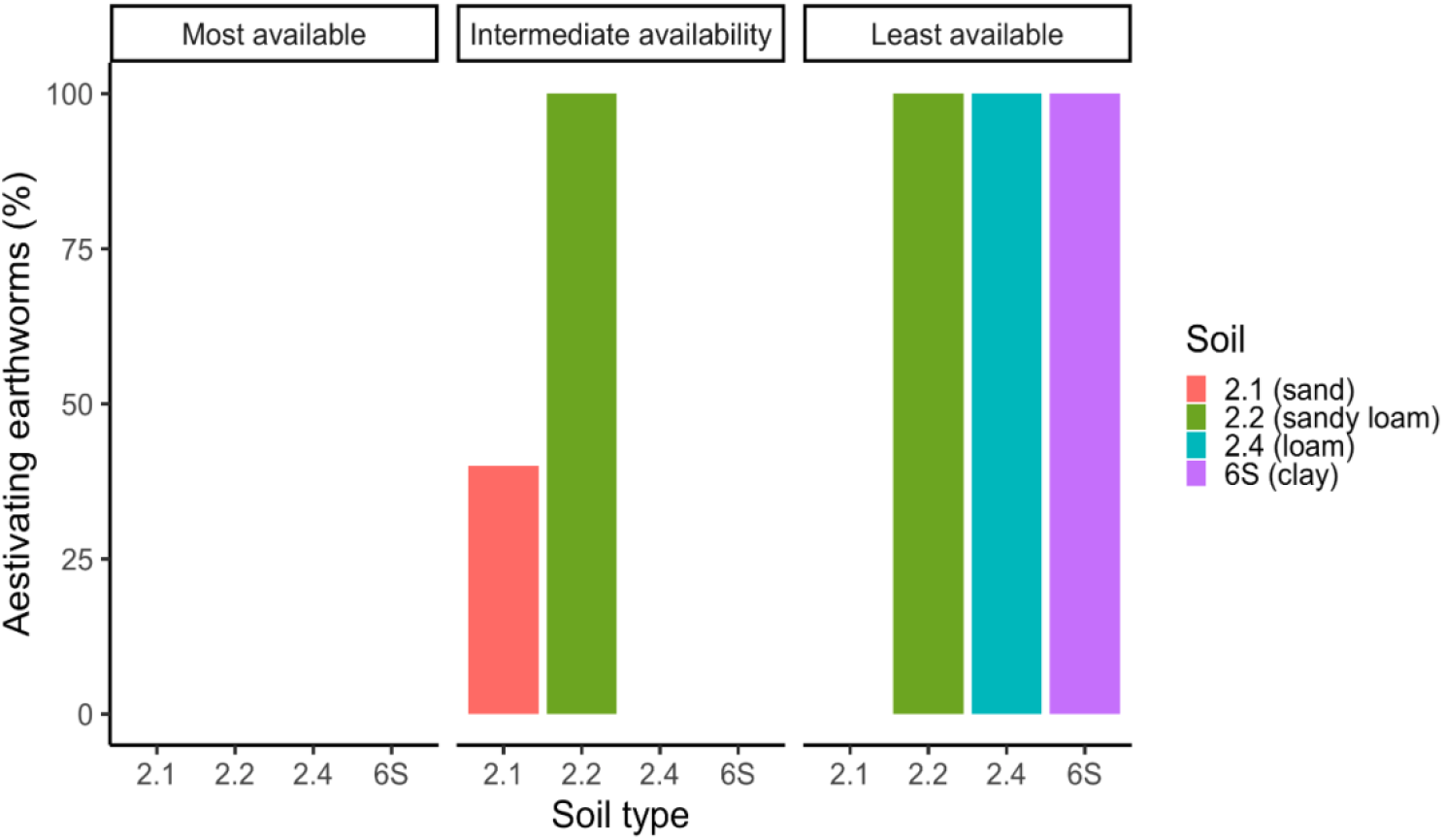
The percentage of *Al. chlorotica* (n = 5) found aestivating at each water potential sampling point (most available = pF ~1.59, intermediate availability = pF ~2.92 and least available = pF ~4.05) in each of the four soil types.

**Figure 3.**
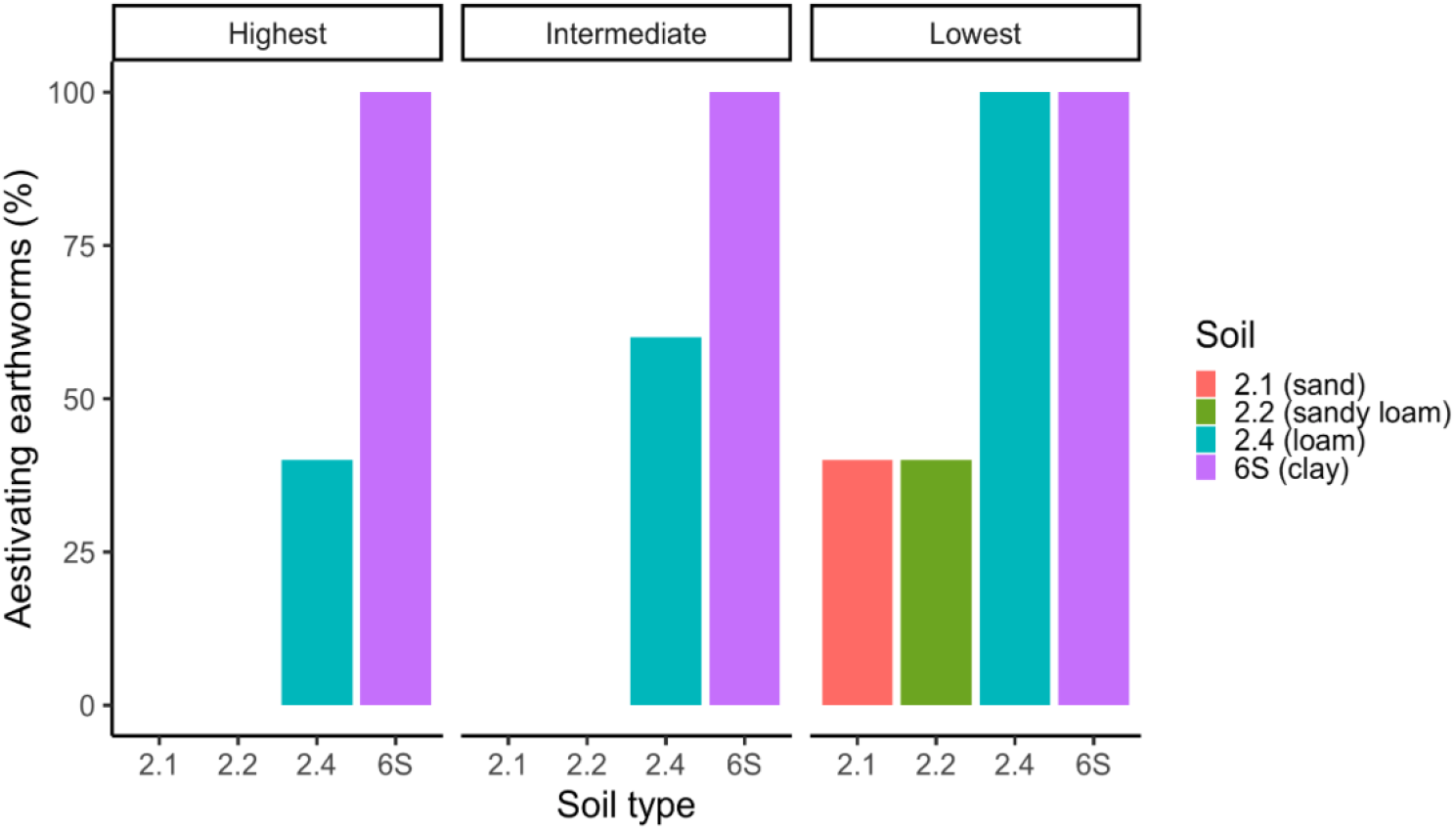
The percentage of *Al. chlorotica* (n = 5) found aestivating at each gravimetric moisture sampling point (highest = ~19.7 wt%, intermediate = ~15.55 wt% and lowest = ~12.39 wt%) in each of the four soil types.

### Change in earthworm mass during hydration

The pre-hydration mass of *Al. chlorotica* differed significantly between active and aestivating individuals (ANOVA: F =205.194, df = 1, 187, p < 0.001), and this relationship varied with soil type (ANOVA: F = 17.66, df = 2, 187, p < 0.001). Prior to hydration, aestivating individuals had significantly lower masses than active earthworms in all soils except the sand (2.1) (Tukey, p < 0.05) (**Fig. 4**). After 24 hours hydration, no significant mass differences remained between earthworms that were initially active or aestivating (Tukey, p > 0.05) (**Fig. 5**). In control conditions, most individuals lost mass during the hydration period, as indicated by points falling below the 1:1 line (**Fig. 6**). In contrast, the direction of mass change under drying conditions depended on behavioural state as aestivating individuals generally gained mass after hydration, while active individuals tended to lose mass (**Fig. 6**).

**Figure 4.**
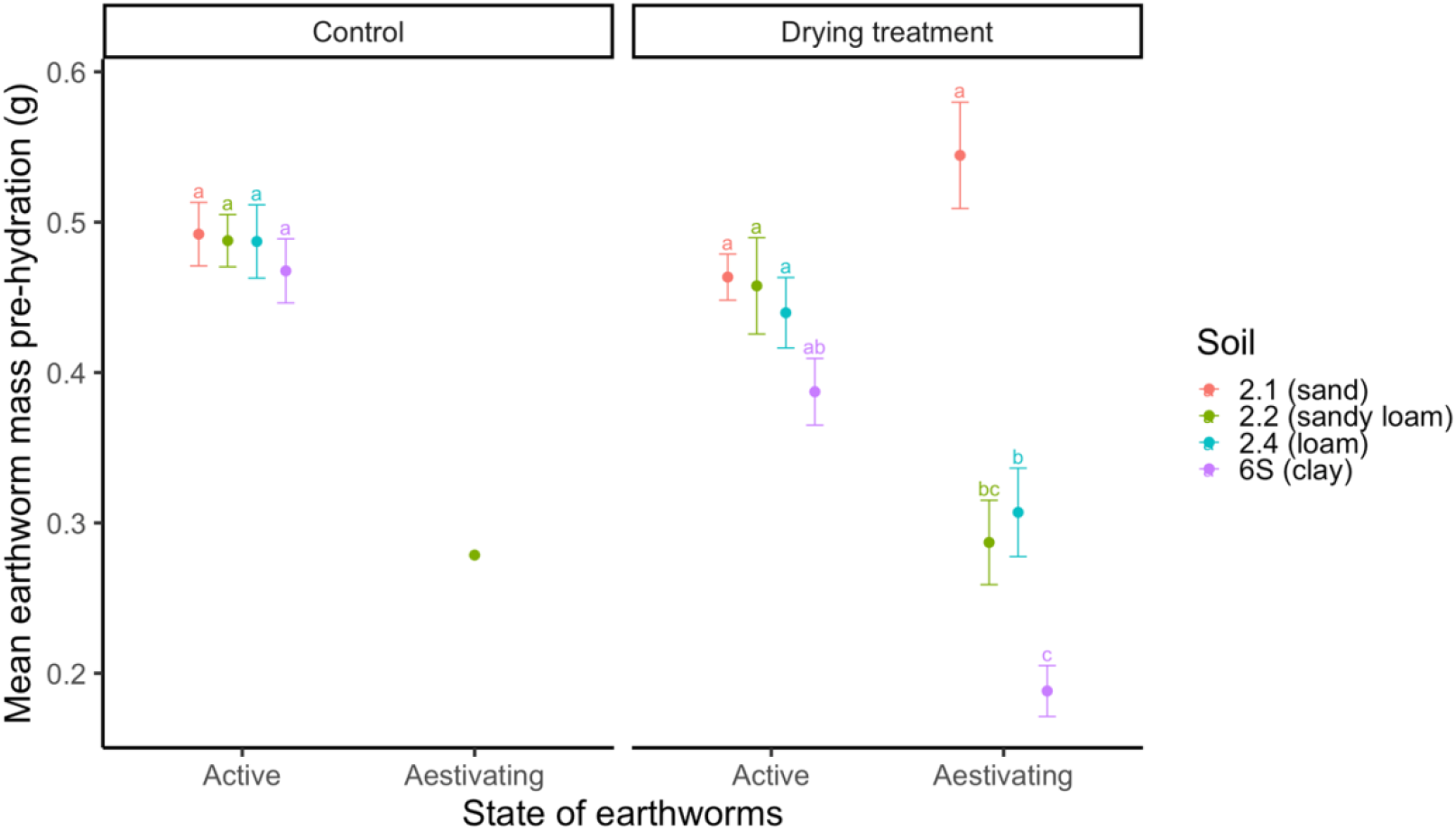
The mean mass of earthworms found active (both in the control and drying treatments) and aestivating in each soil type pre 24 hours hydration. Error bars represent standard error. Treatments with different letters differ to a statistically significant degree (p < 0.05).

**Figure 5.**
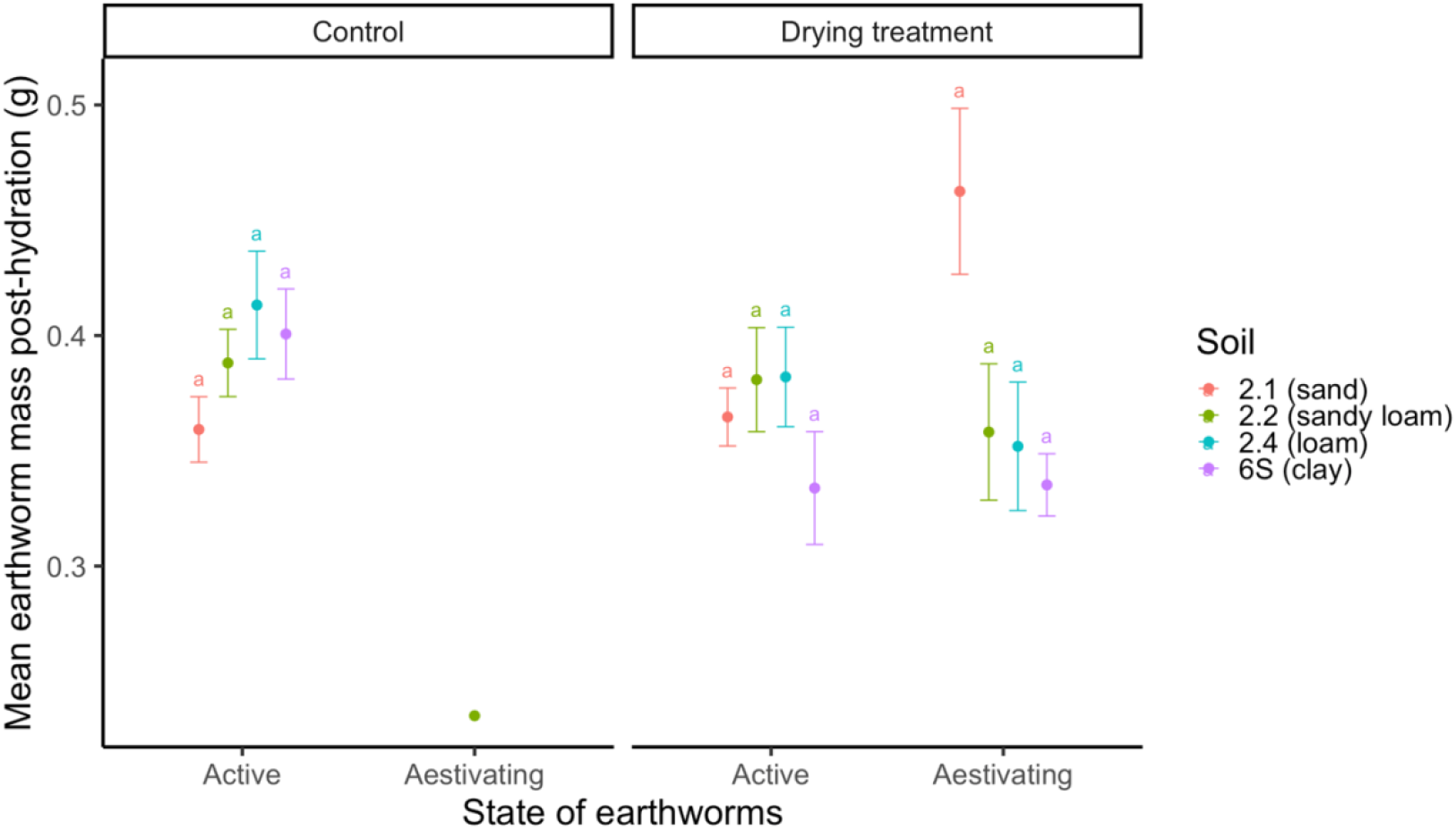
The mean mass of earthworms found active (both in the control and drying treatments) and aestivating in each soil type post 24 hours hydration. Error bars represent standard error. Treatments with the same letter do not differ to a statistically significant degree (p > 0.05).

**Figure 6.**
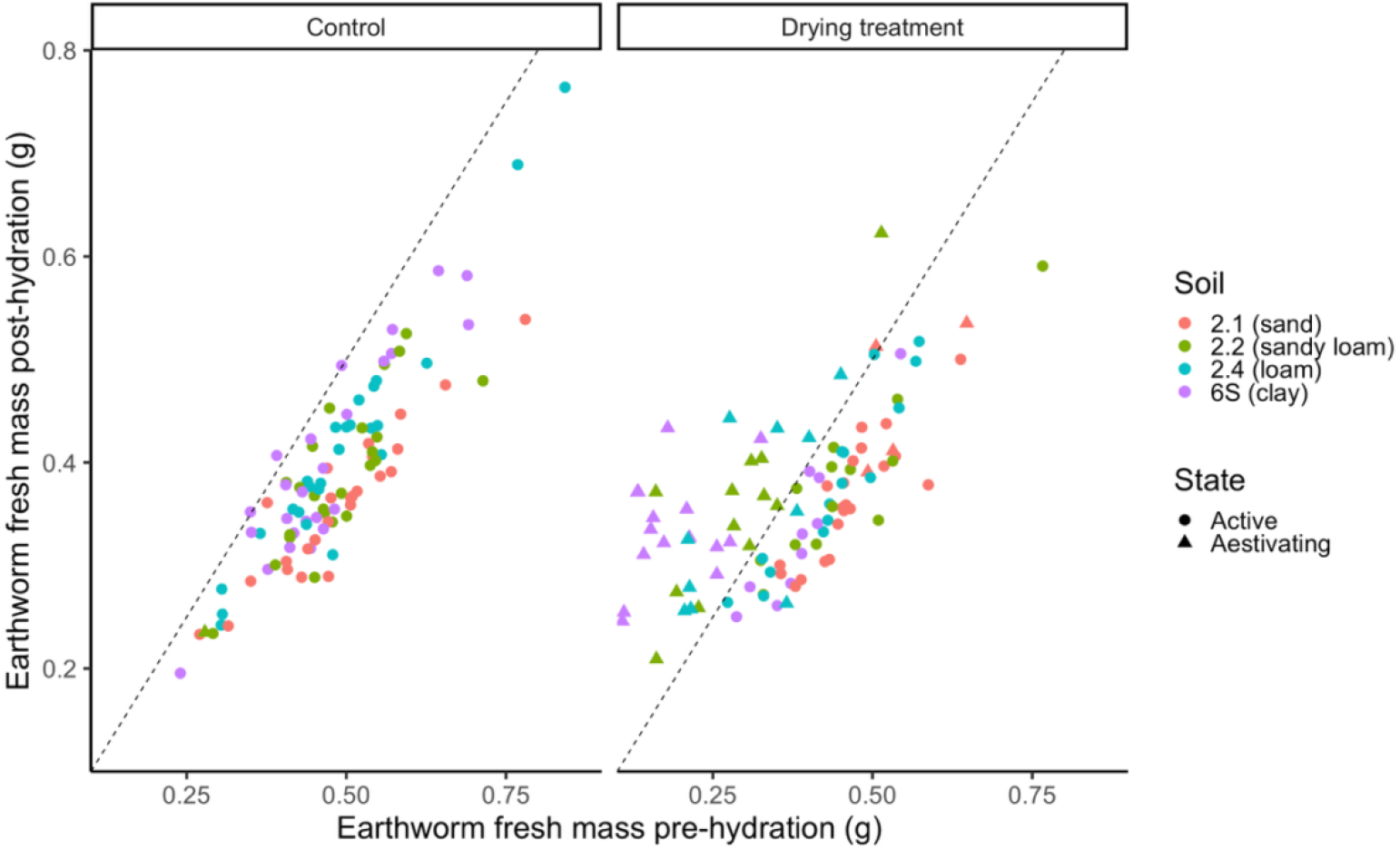
The mass of *Al. chlorotica* pre- and post-exposure to 24 hours hydration, grouped by treatment (constant control or drying conditions). Colours represent each soil type. Shapes represent the state of earthworms on destructive sampling (circles = active, triangles = aestivating). Dashed lines represent no change in mass.

### Influence of water potential (pF) on earthworm mass

There were significant overall main effects of soil type (ANOVA: 12.204, df = 3, 48, p < 0.001) and water potential (pF) (ANOVA: F = 12.423, df = 3, 48, p < 0.001) on the change in *Al. chlorotica* mass measured pre-hydration, with a significant interaction between the two factors (ANOVA: F = 4.228, df = 6, 48, p < 0.01) (**Fig. 7**). At highest water availability (~pF 1.59), all earthworms had increased in mass (by ~36-53 %), and differences among soils were not significant (Tukey, p > 0.05). Similarly, at the intermediate water availability (~pF 2.92), most earthworms gained mass (by ~54-55 %), except those in the sandy loam soil (2.2), which did not differ from their starting mass but differed significantly from earthworms in the other soils (Tukey, p > 0.05). At the lowest water availability (~pF 4.05), earthworms in soils 2.2, 2.4 and 6S had decreased in mass (by ~3 and 26 %) since the start of the experiment, while those in the sand (2.1) still showed an increase (by ~68 %). For those in soils 2.4 and 6S these changes in mass were significantly different to changes in the intermediate water availability (Tukey, p < 0.05).

**Figure 7.**
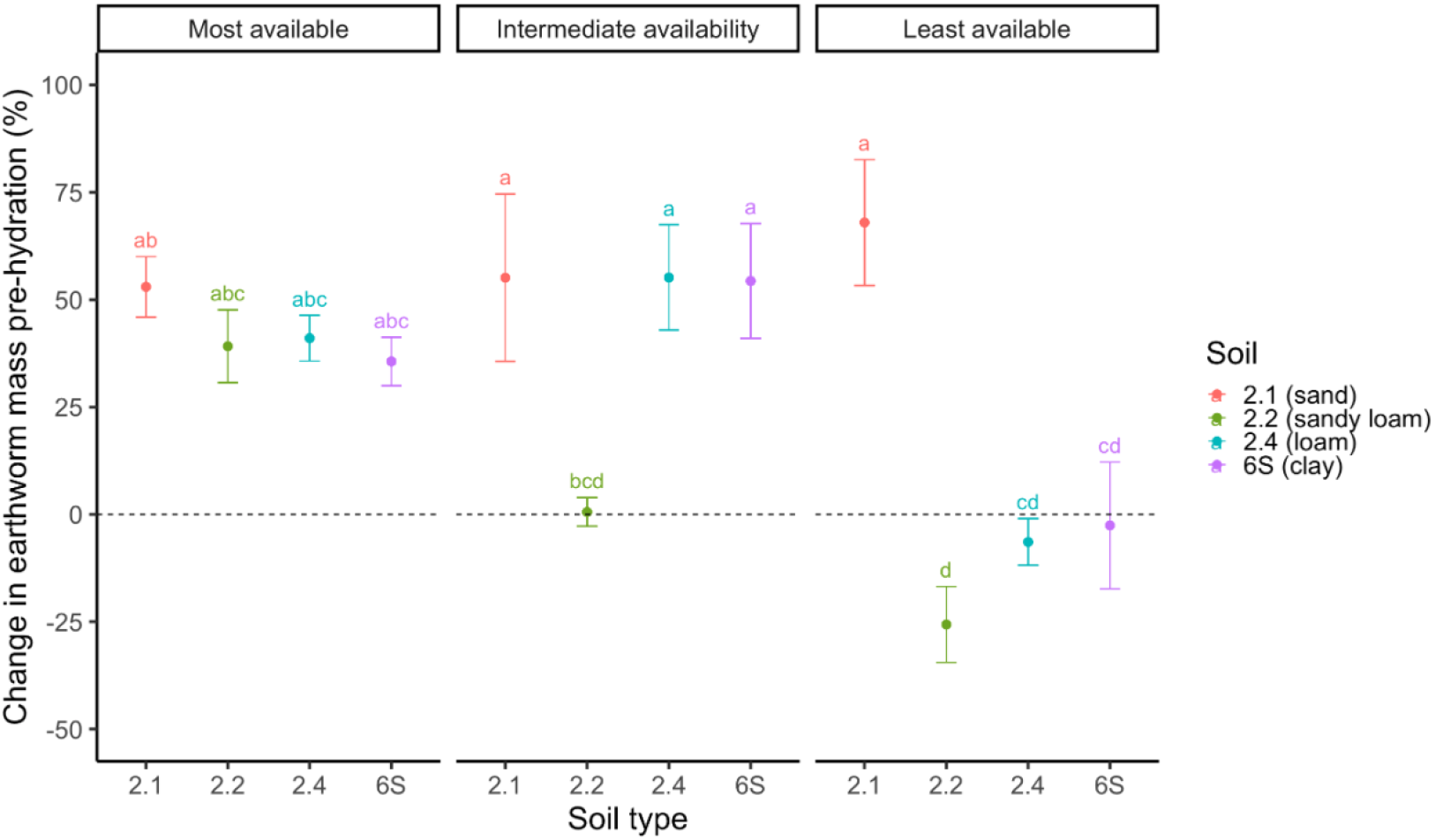
The mean change in earthworm mass (n = 5) relative to starting mass in each soil type at each of the three water potentials (of differing water availability) prior to hydration. Error bars show standard error. Treatments with different letters differ to a statistically significant degree (p < 0.05).

Earthworms in the control conditions had increased in mass (by ~32-120 %) compared to their starting mass at every sampling point (**Fig. S4**). When compared to corresponding control groups, there was a significant interaction between sampling point, soil type and treatment group (ANOVA: F = 2.266, df = 6, 96, p < 0.05). At the highest water availability (pF ~1.59), mass changes in the drying and control groups were similar, but at the lowest water availability (pF ~4.05), mass gains were significantly lower under drying in all soils except 2.1 (Tukey, p < 0.05) (**Fig. S4**).

After 24 hours hydration, all earthworms from the drying treatment had gained mass relative to their initial mass, and there were no significant main effects of water availability (ANOVA: F = 1.008, df = 2, 48, p = 0.372), soil type (ANOVA: F = 1.823, df = 3, 48, p = 0.156), or their interaction (ANOVA: F = 0.847, df = 6, 48, p = 0.541) (**Fig. 8**). However, earthworms previously exposed to drying remained lighter than their controls at the intermediate and least available water potentials (**Fig. S5**). In particular, earthworms in soil 2.2 exposed to the least available water potential had increased by significantly less mass than control earthworms in the same soil (Tukey, p < 0.05).

**Figure 8.**
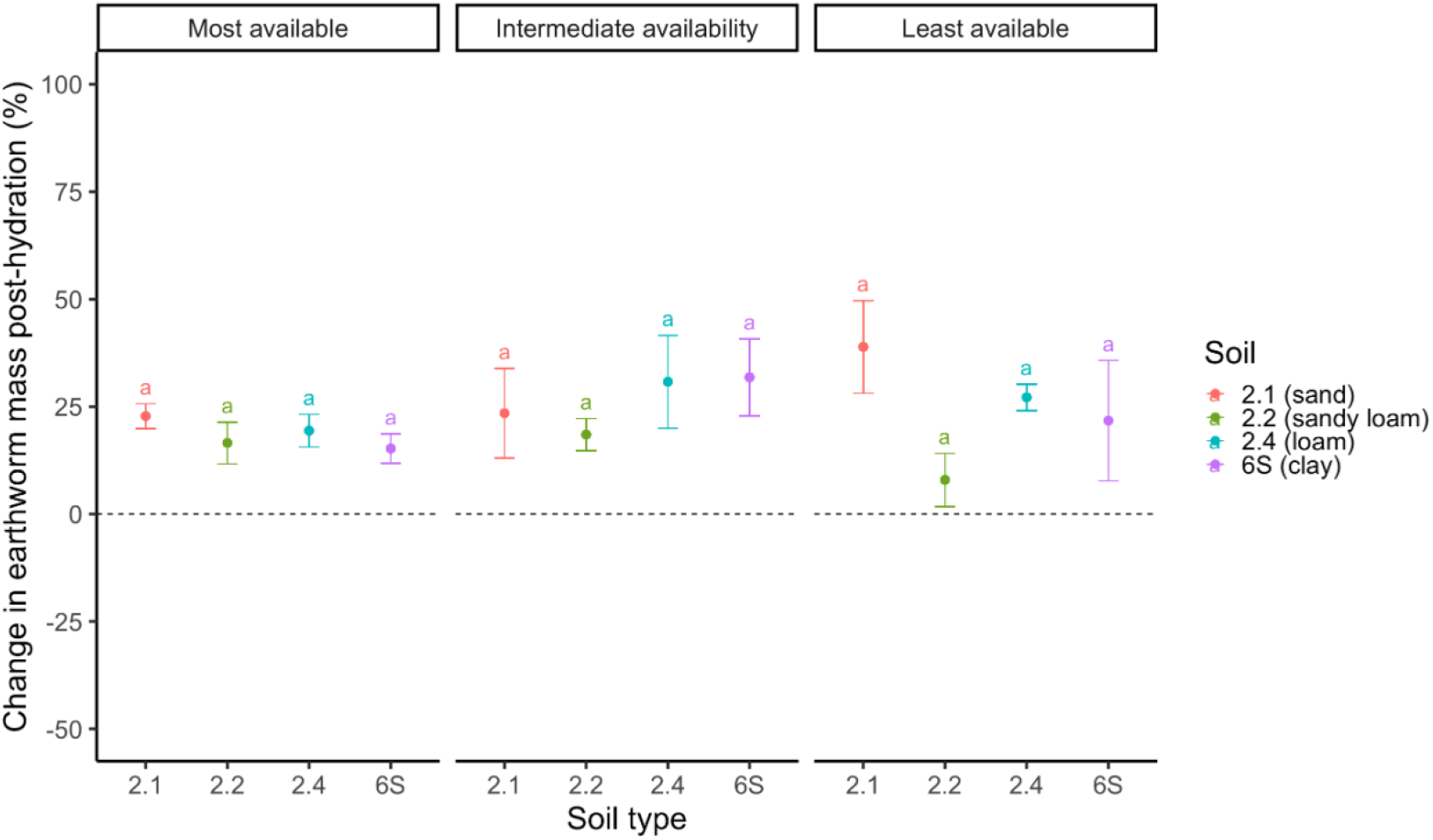
The mean change in earthworm mass (n = 5) relative to starting mass in each soil type at each of the three water potentials (of differing water availability) post-hydration. Error bars show standard error. Treatments with the same letter do not differ to a statistically significant degree (p > 0.05).

### Influence of gravimetric moisture content (wt%) on earthworm mass

Pre-hydration mass changes were significantly affected by soil type (ANOVA: F = 61.587, df = 3, 48, p < 0.001) and gravimetric moisture content (ANOVA: F = 6.581, df = 2, 48, p < 0.01), with a significant interaction between them (ANOVA: F = 4.302, df = 6, 48, p < 0.01) (**Fig. 9**). At the highest moisture content (~19.7 wt%), all earthworms increased in mass (by ~39-53 %) since the start of the experiment, except those in the clay soil (6S), which decreased by ~3 % and differed significantly from those in the sand soil (2.1) (Tukey, p < 0.05). At the intermediate gravimetric moisture content (~15.55 wt%), earthworms in the clay soil (6S) had lost ~45 % of their mass, differing significantly to earthworms in all other soils (Tukey, p < 0.05).

**Figure 9.**
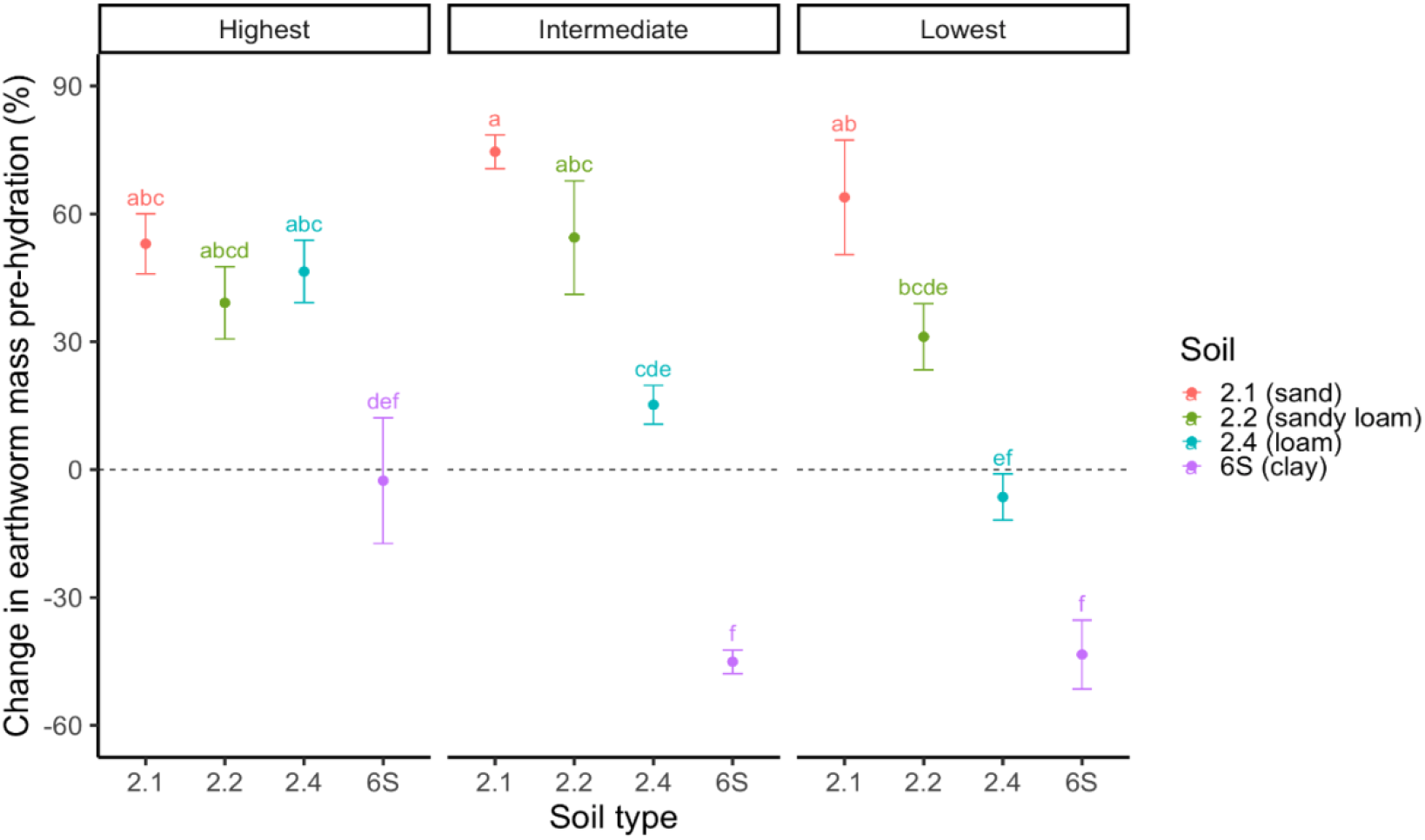
The mean change in earthworm mass (n = 5) relative to starting mass in each soil type at each of the three gravimetric moisture values (of differing water content) pre-hydration. Error bars show standard error. Treatments with different letters differ to a statistically significant degree (p < 0.05).

Earthworms in the other soil types had increased in mass since the start of the experiment, but the increase for those in the loam soil was lower than in the highest moisture content (although not significantly, Tukey p > 0.05). At the lowest moisture content (~12.39 wt%), mass increases persisted only in the sand and sandy loam soils (2.1 and 2.2). Earthworms in the loam and clay soils had lost mass since the start of the experiment (by ~ 6 and ~45 %, respectively), with the change in mass in the loam significantly different than at the highest moisture level (Tukey, p < 0.05).

In the control conditions, all earthworms increased in mass since the start of the experiment (by ~46-99 %) at each sampling point (**Fig. S6**). At the highest moisture content, mass changes in drying and control groups were similar except in the clay soil (6S), where gains were significantly smaller under drying (Tukey, p < 0.05).

Earthworms in the drying clay conditions remained significantly different from their corresponding control in terms of changes in mass at every sampling point (Tukey, p < 0.05). Differences between drying and control treatments became more pronounced at lower moisture contents, and at the lowest level, earthworms in the drying loam soil also differed significantly from their controls (Tukey, p < 0.05).

After 24 hours hydration, all earthworms from the drying treatments gained mass (by ~16-34 %) relative to their initial mass, and no significant effects of soil type (ANOVA: F = 0.325, df = 3, 48, p = 0.807), gravimetric moisture content (ANOVA: F = 0.325, df = 2, 48, p = 0.725), or their interaction (ANOVA: F = 0.414, df = 6, 48, p = 0.866) were detected (**Fig. 10**). Although differences between drying and control groups persisted, mainly in the loam and clay soils, these were not significant (Tukey, p > 0.05).

**Figure 10.**
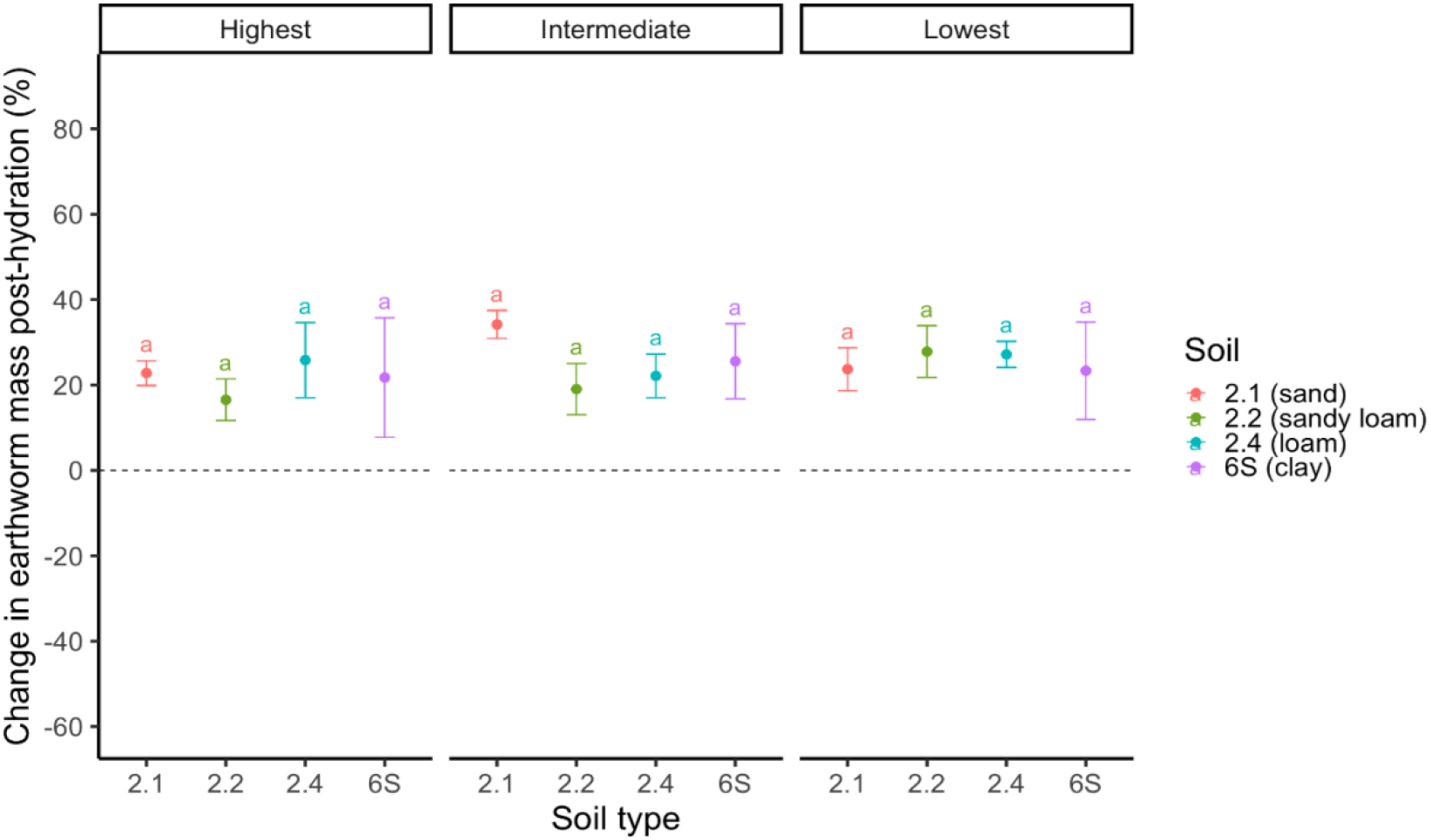
The mean change in earthworm mass (n = 5) relative to starting mass in each soil type at each of the three gravimetric moisture values (of differing water content) post-hydration. Error bars show standard error.

Treatments with different letters differ to a statistically significant degree (p < 0.05).

### Continuous relationships between soil water and earthworm mass

To identify thresholds of mass loss under drying, changes in mass were plotted against continuous measures of water potential and gravimetric moisture (**Fig. 11**). Pre-hydration, both the effect of water potential (Generalised additive model: adjusted R^2^ = 0.708) and gravimetric moisture (Generalised additive model: adjusted R^2^ = 0.712) on earthworm mass change depended strongly on soil type. In the sand soil (2.1), neither water availability (p = 0.661) or gravimetric moisture (p = 0.812) had significant effects, and earthworms consistently gained mass. In contrast, soils 2.2, 2.4 and 6S showed significant nonlinear declines in mass with decreasing water potential (p < 0.001) and gravimetric moisture (p < 0.001). Mass losses began at soil-specific thresholds. Earthworms in the sandy loam (2.2) were most sensitive to changes in water potential, with negative changes in mass observed at higher water availabilities (~pF 2.9) compared to those in the loam (2.4) and clay (6S) soils (in which negative changes were first observed above ~pF 4) (**Fig. 11)**. When expressed in terms of gravimetric moisture content, earthworms in the clay soil were found below their starting mass at just below ~19 wt%, whereas decreases were not observed until ~12 wt% in the loam soil (2.4) and below 10 wt% in the sandy loam (**Fig. 11**).

**Fig 11.**
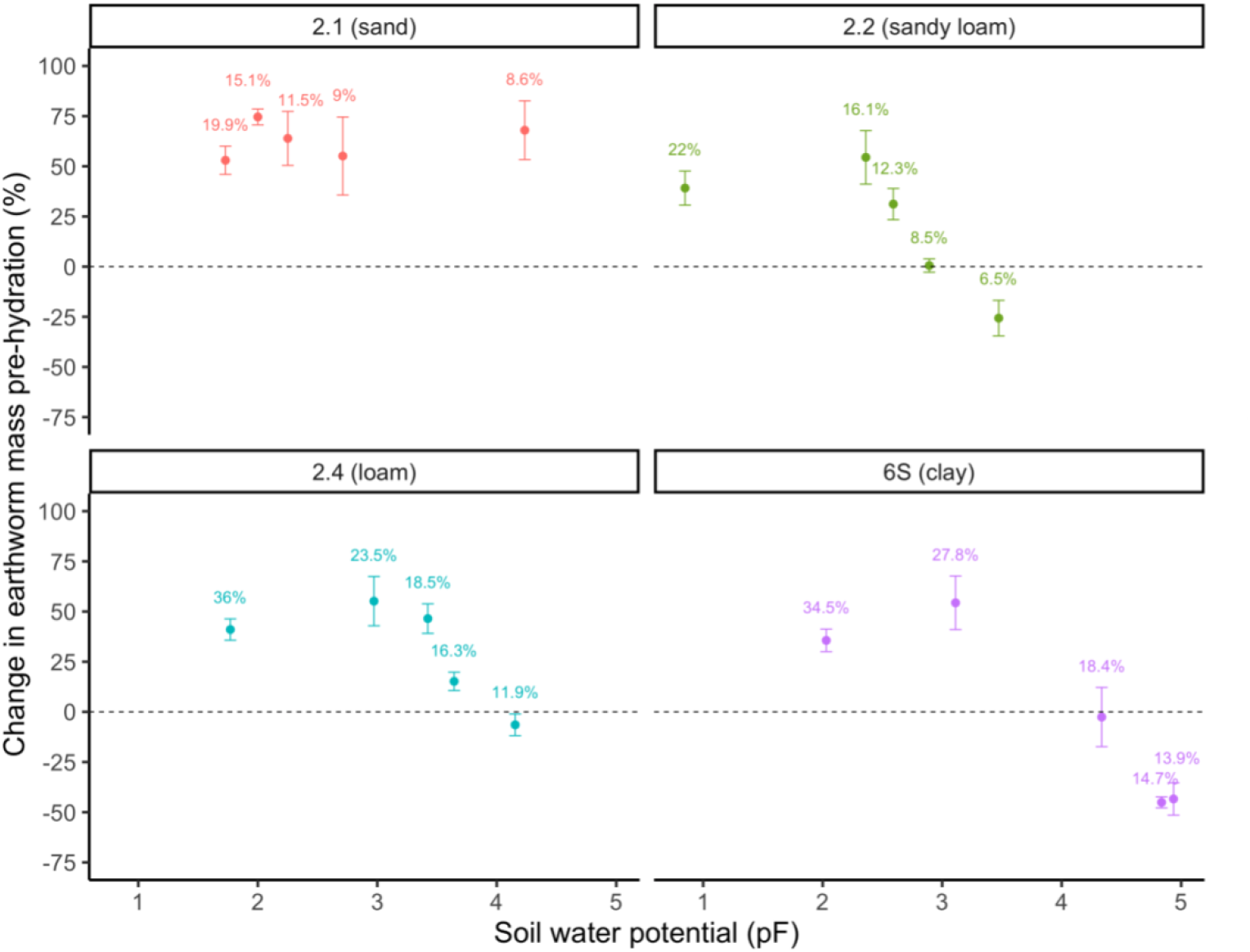
The mean (n = 5) change in *Al. chlorotica* mass relative to their starting mass measured pre-hydration relative to the soil water potential (pF) for each soil type. Error bars show standard error. Percentages above points represent the corresponding gravimetric moisture content for each pF value.

After 24 hours hydration, no significant relationships were detected between water potential (Generalised additive model: adjusted R^2^ = 0.028) or gravimetric moisture content (Generalised additive model: adjusted R^2^ = 0.011) and mass change in all soils, with all individuals greater in mass than at the start of the experiment (**Fig. 12**).

**Fig 12.**
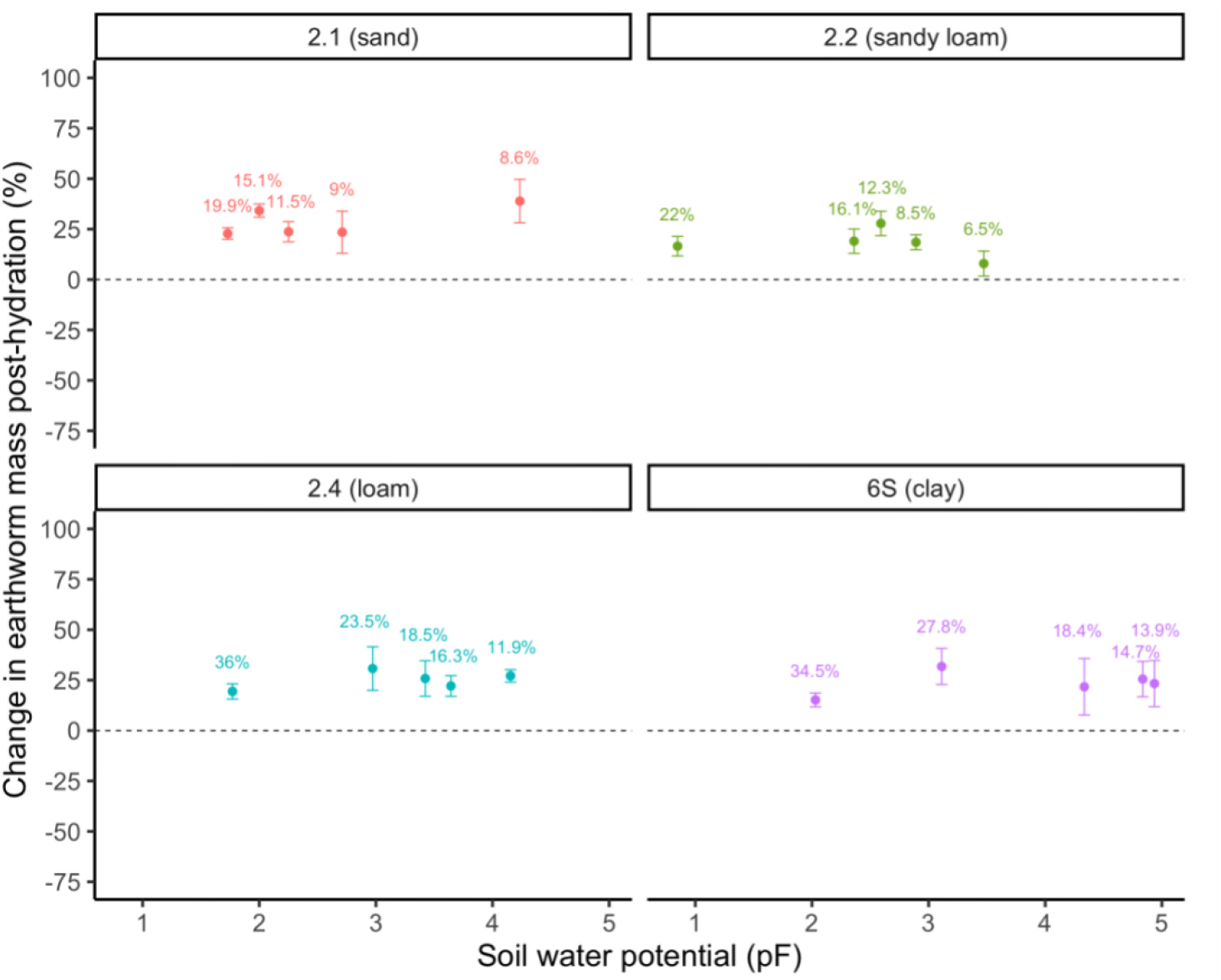
The mean (n = 5) change in *Al. chlorotica* mass relative to their starting mass measured post-hydration relative to the soil water potential (pF) for each soil type. Error bars show standard error. Percentages above points represent the corresponding gravimetric moisture content for each pF value.

## Discussion

In the control conditions (~pF 0.8 to 1.6), nearly all *Allolobophora chlorotica* (99 %) remained active, consistent with Friis *et al*. (2004), who found no *Aporrectodea caliginosa* formed aestivation cells in ‘wet controls’ (18 % moisture, ~pF 1-2). In contrast, under drying, the incidence of aestivation increased significantly with decreasing water potential and varied among soil types. All *Al. chlorotica* were active at the highest water availability (~pF 1.59) and aestivating at the lowest (~pF 4.05), except in the sand (2.1). In the sandier soils (2.1 and 2.2), aestivation was also observed at the intermediate water potential (~pF 2.92). These results align with previous evidence suggesting that lumbricids are generally active only in soils when water is more available than ~pF 3 to 3.3 (100-200 kPa) (Gerard, 1967; Kretzschmar and Bruchou, 1991; Nordström, 1975; Rundgren, 1975; Baker *et al*., 1993). For instance, *Aporrectodea longa* cast production and thus burrowing rates were severely restricted above pF 2.48 (30 kPa) and they induced aestivation at pF 3.22 (Kretzschmar and Bruchou, 1991). In *Ap. caliginosa*, Holmstrup (2001) found all juveniles remained active at water potentials of ~pF 2.08 (12 kPa) but entered aestivation above ~pF 2.3 (20 kPa), with a steep increase between pF 2.3 to 2.48 (20 to 30 kPa) and almost all aestivating at pF 2.6 (40 kPa). Later studies confirmed that adult *Ap. caliginosa* also form aestivation chambers in soils of 30 kPa (Bayley *et al*., 2010), suggesting that these lower thresholds are not a result of earthworm maturity. Therefore, these different thresholds likely represent species-specific differences in drought tolerance which may reflect physiological differences.

Water availability, soil type, and their interaction significantly affected changes in *Al. chlorotica* mass on immediate removal from the soil. At the most available (~pF 1.59) and intermediate (~pF 2.92) water potentials, earthworms generally increased in mass relative to their starting mass, although gains in the sandy loam (2.2) were negligible and significantly lower than their corresponding control group. At the least available water potential (~pF 4.05), earthworms lost up to ~26 % of their initial mass, except those in the sand. Similarly, Kretzschmar and Bruchou (1991) found *Aporrectodea longa* lost as much as 60 % of their initial weight when exposed to pF 3.5 and 4.3. Water potentials below pF 2.78 had little effect on *Ap. longa* mass, although mass did still fluctuate, which they suggest is likely due to losses from casting and gains from ingestion (Kretzschmar and Bruchou, 1991). In contrast, *Aporrectodea caliginosa* mass declined at water potentials greater than ~pF 2.08 (12 kPa) (Holmstrup, 2001). These findings support species-specific differences in tolerance. These responses may relate to osmotic regulation as species differ in their osmotic pressures. It has been suggested that earthworms generally lose body water through osmosis when the soil water potential increases above ~pF 3.6 (~400 kPa) as the osmotic pressure of their body fluids becomes higher than that of the surrounding soil (Bayley *et al*., 2010; McDaniel *et al*., 2013a; Holmstrup *et al*., 2016). If aestivation serves as a precautionary mechanism before critical desiccation (Holmstrup *et al*., 2016), aestivation should be induced at water potentials below that of their body fluid osmolality. Indeed, *Al. chlorotica* aestivated at ~pF 2.92 in the sand and sandy loam soils, whereas those in the loam and clay tolerated lower water availabilities and only aestivated at ~pF 4.05. This supports findings of a significant positive correlation between the percentage of clay and the water potential (pF value) at which the distribution and thus behaviour (avoidance) of *Aporrectodea trapezoides* did not diverge from the controls (Doube and Styan, 1996).

When expressed as gravimetric water content, both earthworm activity and mass change were significantly influenced by soil moisture, though variation was explained more strongly by soil type. In the sand and sandy loam, aestivation was absent until the lowest sampling point (~12.39 wt%), where over half of the individuals remained active. These responses are consistent with our previous observations of *Al. chlorotica* remaining active at 18 wt% aestivating at 13 wt% (unpublished results), and with studies of *Hormogaster elisae*, which rarely aestivated at 20 % moisture but did so extensively at 10 % moisture, particularly in spring and summer (Díaz Cosín *et al*., 2006). Moreover, in the present study, earthworms in the sand and sandy loam remained greater than their initial mass even at the lowest of the three gravimetric moisture sampling points (~12.39 wt%). In contrast, in the loam and clay soils, aestivation occurred even at the highest of the three gravimetric sampling points (~19.9 wt%), with 100 % aestivation and a ~3 % mass loss for those in the clay soil. This pattern supports the prediction that aestivation and desiccation occur at higher moisture contents in clay-rich soils because water is held more tightly and is less biologically available at a given gravimetric content. These contrasts in responses highlight the limitations of using gravimetric moisture alone as soils differ in water-retention characteristics, and equal water contents can correspond to very different water potentials.

The sandy loam (2.2) used here resembled soils in Díaz Cosín *et al*. (2006), explaining the similar behavioural thresholds, whereas at ~19.9 wt% the loam and clay already had reduced water availability (>pF 3 and >pF 4, respectively). Similarly, Doube and Styan (1996) reported that the threshold at which *Ap. trapezoides* avoided soils was influenced by an interaction between water content and soil type and ranged from 10 wt% (15 kPa, pF 2.2) in sandy loam, 12 wt% (25 kPa, pF 2.4) in loam and 20 wt% (300 kPa, pF 3.4) in clay. However, Doube and Styan (1996) found water potential was a more important factor than bulk content in determining the behaviour of *Aporrectodea rosea* as they moved out of soil at pF 3.4 (300 kPa), irrespective of soil type or gravimetric moisture content which ranged from 7 - 20 wt%. Therefore, gravimetric thresholds are soil-dependent, and both water potential and soil texture are necessary to predict species responses under drought.

Despite considerable mass losses of *Al. chlorotica* under drying conditions, no mortality occurred. Similarly, Holmstrup (2001) found *Ap. caliginosa* survived exposure to 14-day drought periods with water potentials as low as pF 3.53 (Holmstrup, 2001). Moreover, there is evidence that *Al. chlorotica* can survive and recover from losses of up to 75 % of their body water (Roots, 1956). After just 24 hours of hydration, all *Al. chlorotica* had gained mass since the start of the experiment (from 0.291 ± 0.074 g to 0.376 ± 0.087 g), irrespective of the soil type and degree of drying they had experienced. Consequently, although aestivating individuals were significantly lower in mass than those found active (except for in sand) pre-hydration, they regained water rapidly, while active earthworms lost mass via gut evacuation during hydration, resulting in no significant differences post-hydration. This is consistent with McDaniel *et al*. (2013a) who found that *Ap. caliginosa* exposed to 1-, 2-, or 3-week cycles of drought stress were able to rehydrate when returned to higher moisture conditions, with no lasting negative effects on their biomass. Similarly, Díaz Cosín *et al*. (2006) found *H. elisae* recovered their initial body mass once placed into soil with a higher moisture content (20 wt%), although over a longer period of around one week (6.5 ± 3.6 days). Such rapid recoveries of mass support the view that mass losses largely reflect water loss through urine production, gut evacuation and mucus secretion in the construction of aestivation chambers (Holmstrup, 2001).

Although changes in mass were positive and less marked overall after hydration, some differences in final mass remained between drying and control earthworms, particularly in those subjected to the greatest degree of drying. Assuming *Al. chlorotica* were fully hydrated after 24 hours on wet filter paper, this suggests that some differences were a result of lower tissue mass. Earthworms in the control conditions remained active, continued feeding and gained mass throughout (**Fig. S8, 9**), suggesting that food was not a limiting factor restricting growth. Instead, such reductions in mass may reflect suspended feeding (Reinecke and Reinecke, 2007) and the regression of secondary sexual structures such as the clitellum (**Fig. S10**) characteristic of aestivation in earthworms (Olive and Clark, 1978; Jiménez *et al*., 2000). We also observed constricted caudal segments which suggest selective dehydration and reduction of non-vital tissues. Jiang *et al*. (2023) state that some tissue loss during aestivation may be a result of adipose tissue being broken down to produce adenosine triphosphate (ATP), while other organs such as those involved in digestion may lose mass through lack of use. In earthworms, some tissue loss may be compensated for as undifferentiated stem cells allow for regeneration of localised body segments by developing blastema into new tissue (Christyraj *et al*., 2025).

Interestingly, *Al. chlorotica* in the sand (2.1) gained mass irrespective of the degree of drying and were rarely observed aestivating, even at the least available water potential (~pF 4.05). In contrast, aestivation was common in the loam and clay soils (2.4 and 6S) which had considerably lower sand contents (33.1 % and 22.4 %, respectively, versus 88.2 % for 2.1). The loose texture and lower density of the sand likely permitted movement and feeding even as it dried, while the loam and clay soils hardened as they dried, increased mechanical resistance which constrains burrowing and increases the energetic costs of movement (Arrazola-Vasquez *et al*., 2022). This higher energetic cost combined with the reduction in soil water likely resulted in the onset of aestivation. Moreover, although absolute water loss (g/day) was similar across soils (~1 g/day, **Fig. S11**), the rate of water potential change differed markedly. The sand had a steep water retention curve at low pF values meaning that water was easily lost from the soil until only the tightly held water remained (**Fig. 1**). Consequently, the sand soil (2.1) was the quickest to reach the driest water content (pF 4.05), only 19 days after the most available water potential (pF 1.59), whereas the sandy loam, loam and clay soils took 39, 40 and 32 days, respectively (**Table S1**). This meant that earthworms in the sand soil experienced a shorter duration of physiologically stressful conditions, likely allowing them to remain active. Previous studies show that the proportion of *Ap. caliginosa* aestivating increases with drought duration (McDaniel *et al*., 2013a). Similarly, Díaz Cosín *et al*. (2006) found that while some *H. elisae* maintained an active state after 2 weeks at 10 wt% moisture, 4 months at 10 % moisture resulted in 100 % aestivating. In our experiment, soils were gradually dried rather than kept at constant water levels, thus future work is needed to investigate whether longer exposures could induce aestivation in *Al. chlorotica* even at higher water potentials.

Moreover, aestivation chamber characteristics also varied with soil type. Chambers in sandy soils were fragile and poorly sealed, whereas those in the clay-rich soils were more durable and remained intact during destructive sampling. Similarly, Bayley *et al*. (2010) observed that a sandy loam soil (35 % coarse sand, 45 % fine sand, 9.4 % silt, 8.9 % clay) contained aestivation chambers which broke apart on handling, while those in a slightly less sandy soil (38.4 % coarse sand, 23.6 % fine sand, 22.3 % silt, 13 % clay) were more suitable for handling and extraction. This suggests that soil texture constrains chamber construction and thereby the feasibility of aestivation. Generally, soils with > 80 % sand are considered unsuitable for earthworms and constrain their distribution and activity, due to the abrasive nature of sand particles (Booth *et al*., 2000; El-Duweini and Ghabbour, 1965). However, some studies suggest coarse sand can support earthworms and increase digestive abilities by aiding in the grinding of plant residues in the gut (Eisenhauer *et al*., 2009) although, the effect of sand content was negative for *Octolasion tyrtaeum*, which increased in abundance and biomass with decreasing sand content, while there was no effect on *Ap. caliginosa* (Eisenhauer *et al*., 2009). Thus, the suitability of sandy versus clay soils depends on species traits and ecological strategy, as coarser soils may facilitate continued activity under short-term drought, while finer soils better support aestivation through chamber stability and moisture retention.

## Conclusion

Overall, this study demonstrates that the thresholds inducing aestivation in *Allolobophora chlorotica* depend not only on the degree of drying but also on soil type. Across soils, earthworm activity and changes in mass were most consistent at equal water availabilities, whereas responses diverged at equal gravimetric moisture contents, with more of the variation explained by soil type. This suggests that water potential is a stronger predictor of behaviour than bulk water content. Although, soil texture modulated these responses as aestivation and mass loss occurred at lower pF values in sandier soils. *Al. chlorotica* lost up to 45 % of their initial mass under the driest conditions, and those that aestivated tended to be significantly smaller than those that remained active. However, earthworms rehydrated rapidly upon rewetting, regaining or exceeding their starting mass within 24 hours. This recovery highlights the effectiveness of aestivation and suggests that mass losses are transient and mostly reflect body water loss. Despite this, residual mass differences and regressed sexual structures in some individuals suggest that prolonged or repeated drying may impose physiological costs, potentially reducing reproductive output. Overall, *Al. chlorotica* are incredibly resilient to desiccation, but their responses to drought are shaped by complex interactions between soil texture and water availability. These factors determine not only their physiological responses and changes in mass but also the restrictions imposed on their movement, and their ability to construct aestivation chambers, which is notably more successful in soils with a higher clay content. Further studies across lumbricid species and soil types are needed to refine predictive thresholds for drought-induced aestivation, improving our understanding of how soil fauna activity and ecosystem functions may shift under increasing drought frequency with climate change.

## Acknowledgements

The study was supported by an Adapting to the Challenges of a Changing Environment Doctoral Training Partnership studentship (NE/S00713X/1).

## Author contributions

Conceptualisation, RVAB, PJW, MEH; Investigation, RVAB; Methodology, RVAB, PJW, MEH; Supervision, PJW, MEH; Formal analysis, RVAB; Writing – original draft, RVAB; Writing – review and editing, RVAB, PJW, MEH.

