## Supplementary materials for "The response of *Allolobophora chlorotica* to drought stress in four soils"

**Figure S1.** The relative proportion (%) of clay, silt and sand particles within the four soils: 2.1 (blue), 2.2 (green), 2.4 (purple) and 6S (red). Based on data of mean particle size distribution (The chemical and physical characteristics of standard soils, LUFA Speyer 2022).

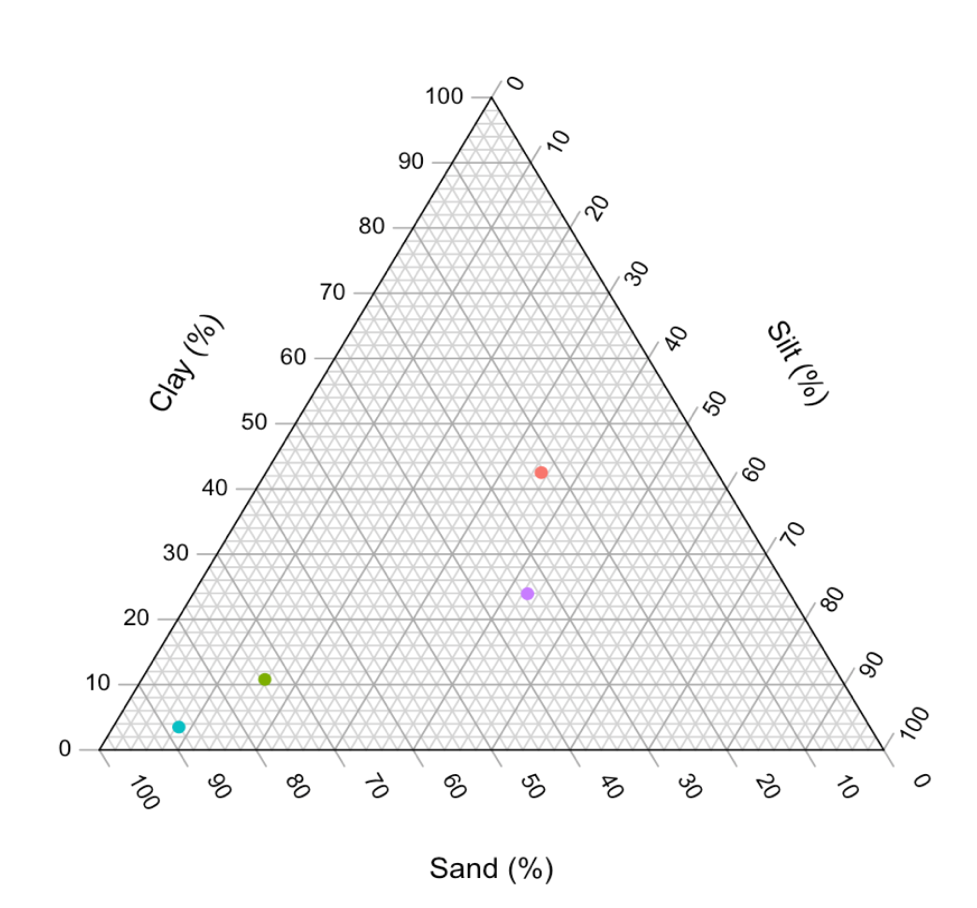

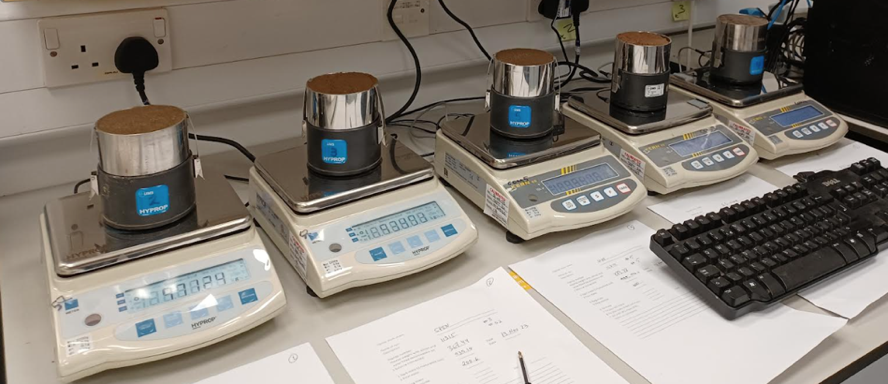

**Figure** **S2**. Hyprop devices containing a saturated soil core placed on a balance, used to determine the water retention curves of each soil type.

HYPROP method:

Each soil core (250 ml) was packed with one of the four soil types, a nonwoven cloth was placed on one side along with a saturation plate before the core was gently placed into a tub filled with deionised water up to 5 mm below the sample rim and left overnight to fully saturate through capillary action until the surface was shiny. The tensio-shafts have a porous ceramic tip and a shaft which was filled with water which was degassed by vacuum to avoid air bubbles. The sensor unit was also filled with degassed water. Each tensio-shaft was gently screwed into the sensor unit and the pressure was monitored to ensure it was increasing but not exceeding 200 kPa to avoid damaging the pressure sensor. A silicone gasket was added to avoid soil entering the sensor unit and several function checks (speed of response, zero point etc.) were carried out to ensure the pressure was responding as intended. Two holes were made in each soil core using an auger and the sensor unit was inverted over the soil core so that the tensio-shafts could be carefully inserted into the holes making sure not to compress the soil. The soil core was upturned, saturation plate and cloth removed and then the soil sample was fixed to the sensor unit using clips and placed onto a balance. The balance was connected to a computer with the HYPROP software and measurements commenced. Recording was terminated when the soil sample had reached the air entry phase whereby the tension value drops abruptly to zero as air enters the ceramic tips. The soil core was removed from the sensor unit and oven dried at 105 °C for 24 hours to determine the dry weight of the soil sample.

**Figure S3**. Mean initial fresh mass of *Al. chlorotica* (n = 25) when weighed prior to the start of the experiment for each of the four soil types, grouped according to treatment group for those that would be subjected to either constant control or drying conditions. Error bars show standard error. Treatments with the same letter do not differ to a statistically significant degree (p > 0.05).

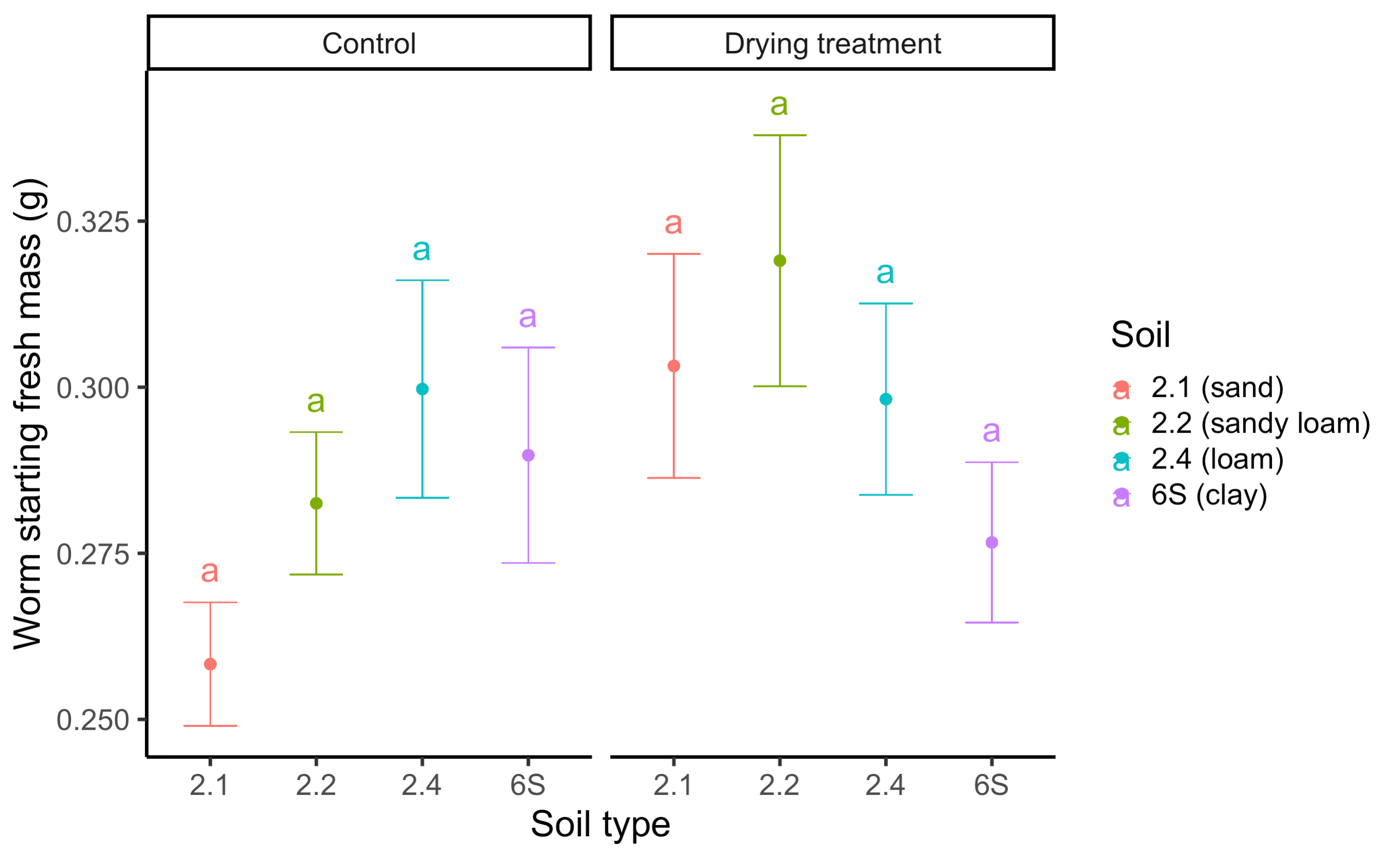

**Figure S4**. The mean change in earthworm mass (n = 5) relative to starting mass at each of the three water potentials (wet, medium and dry) for each soil type prior to hydration. Earthworms in the constant control conditions (blue) were sampled at the same time and those in the drying soils (red). Error bars show standard error. Treatments with different letters differ to a statistically significant degree (p < 0.05).

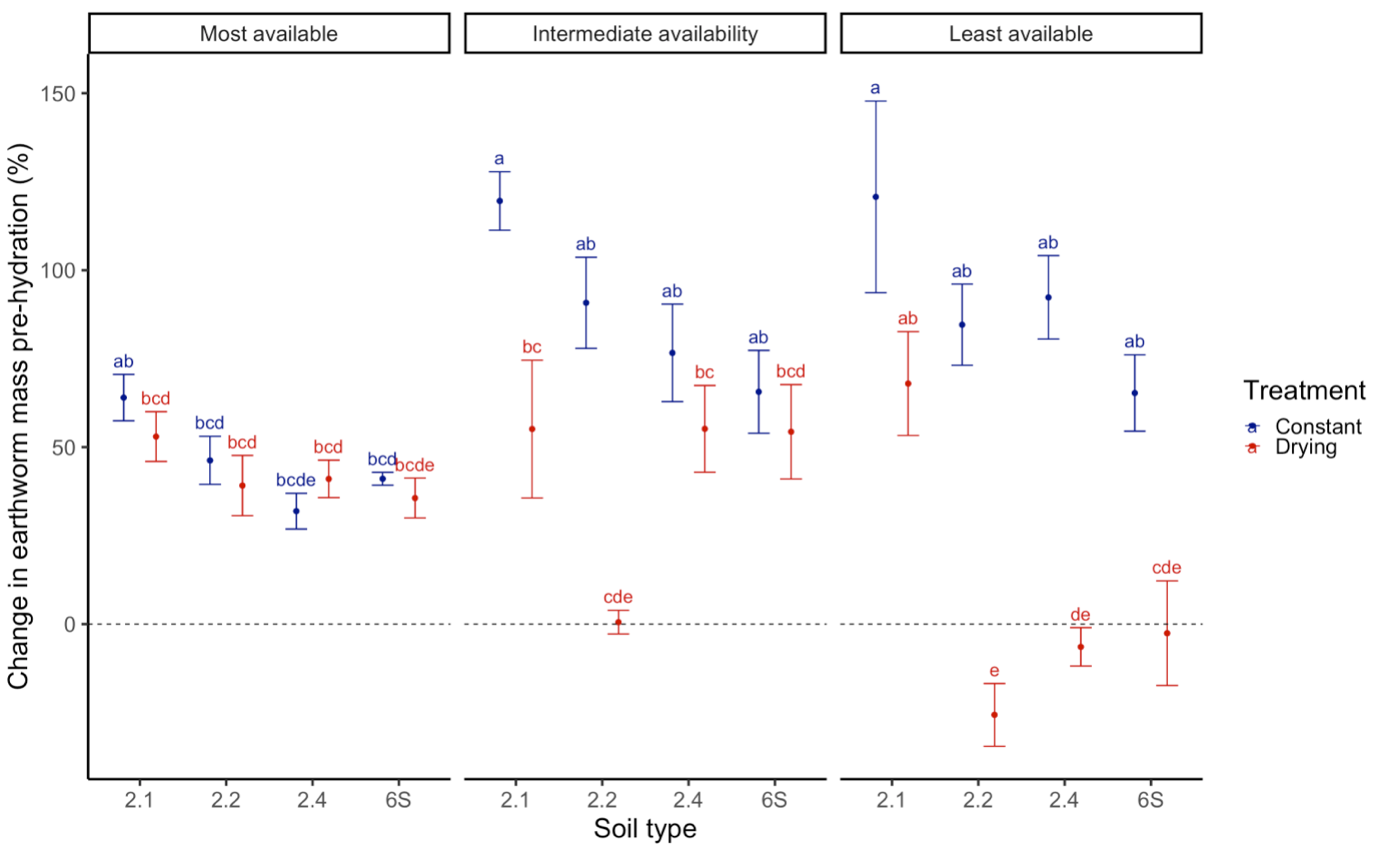

**Figure S5**. The mean change in earthworm mass (n = 5) relative to starting mass at each of the three water potentials (wet, medium and dry) for each soil type after 24 hours hydration. Blue = constant control conditions, red = drying soils. Error bars show standard error. Treatments with different letters differ to a statistically significant degree (p < 0.05).

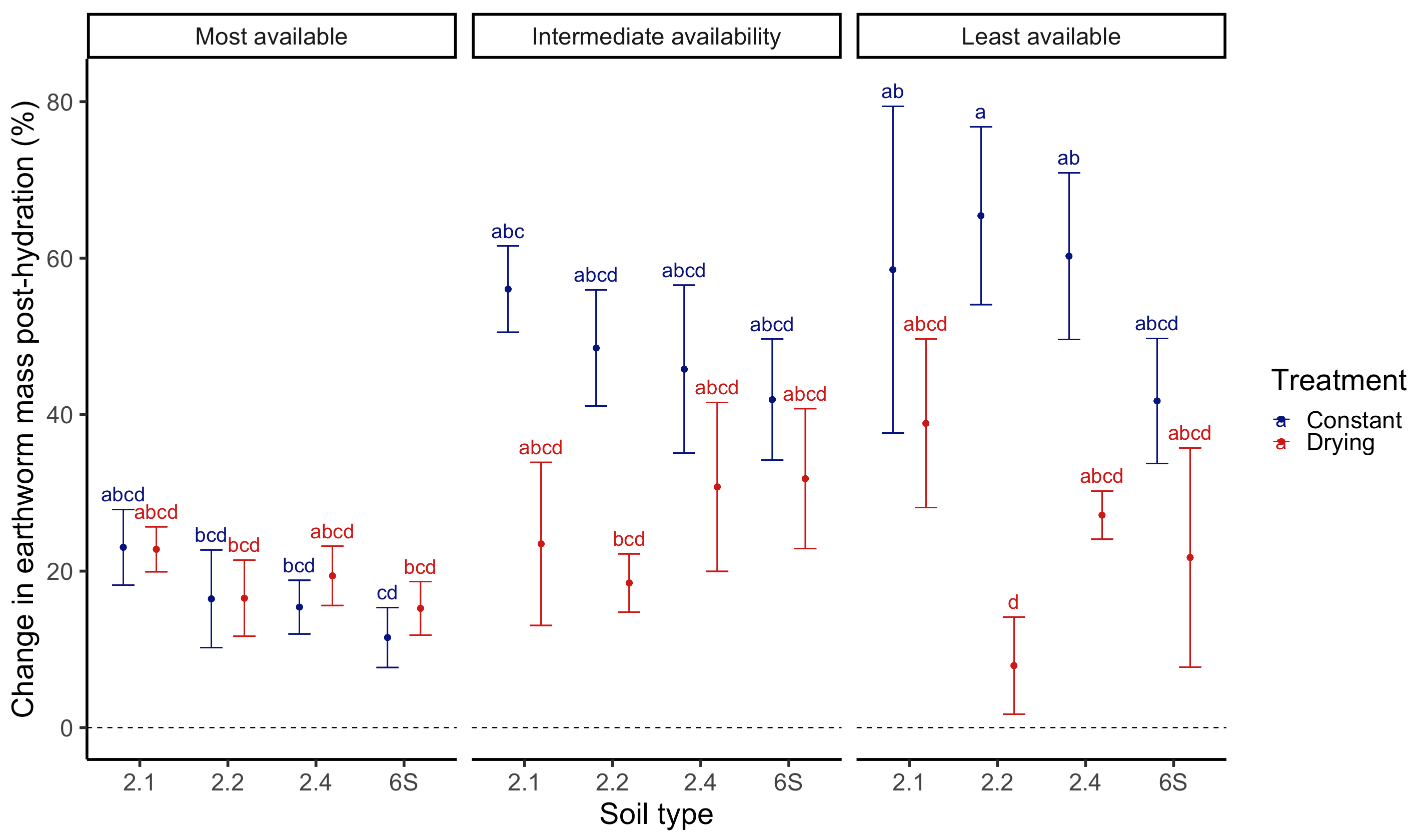

**Figure** **S6**. The mean change in earthworm mass (n = 5) relative to starting mass at each of the three gravimetric moisture sampling points (highest, intermediate and lowest water contents) for each soil type pre-hydration. Earthworms in the constant control conditions (blue) were sampled at the same time and those in the drying soils (red). Error bars show standard error. Treatments with different letters differ to a statistically significant degree (p < 0.05).

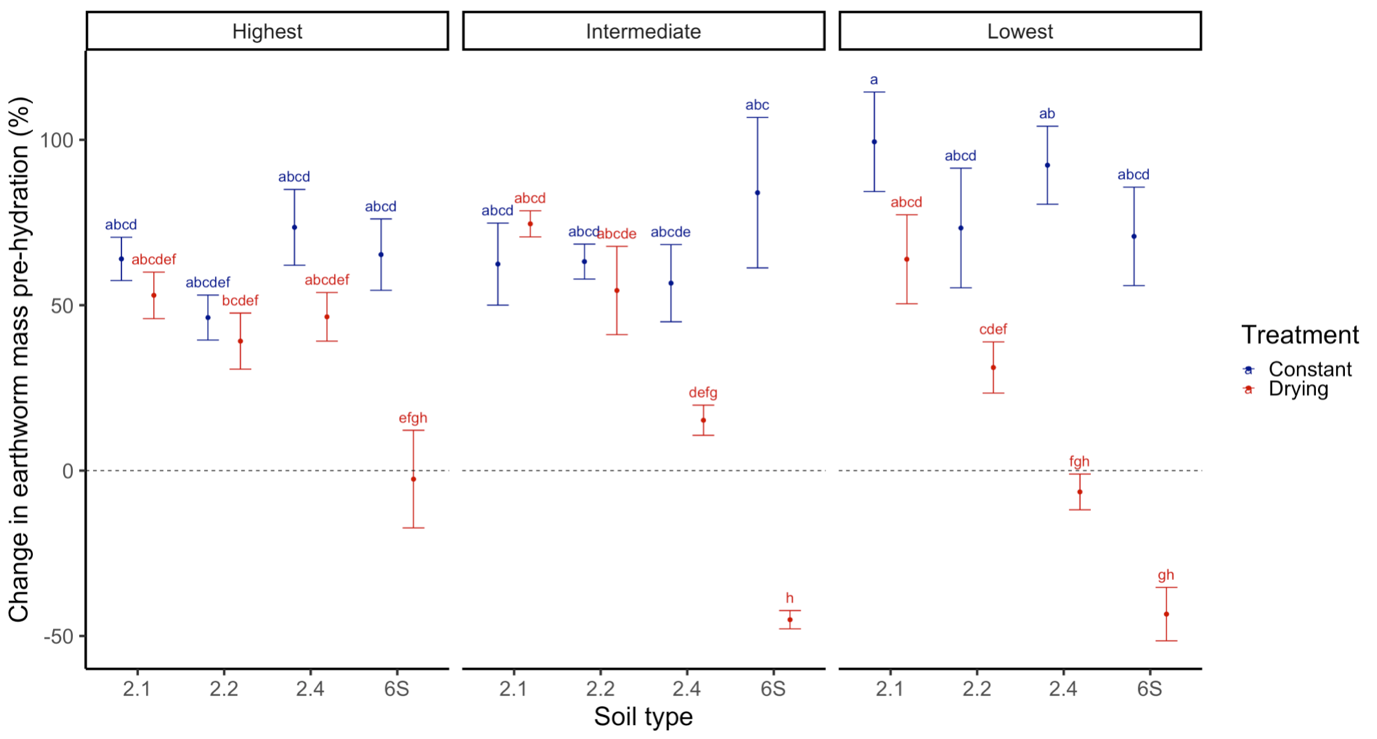

**Figure S7**. The mean change in earthworm mass (n = 5) relative to starting mass at each of the three gravimetric moisture sampling points (highest, intermediate and lowest water contents) for each soil type after 24 hours hydration. Earthworms in the constant control conditions (blue) were sampled at the same time and those in the drying soils (red). Error bars show standard error. Treatments with different letters differ to a statistically significant degree (p < 0.05).

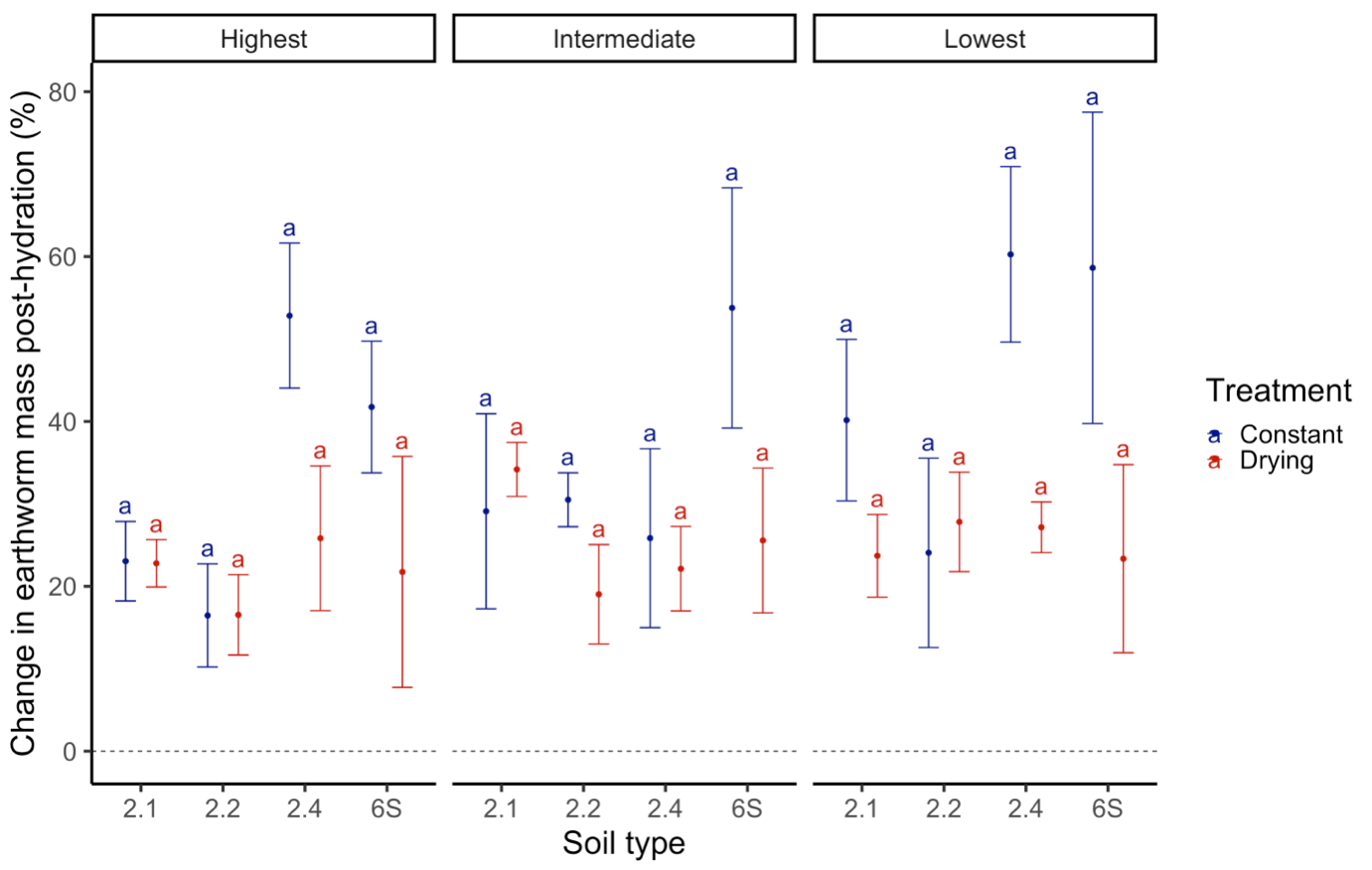

**Figure S8**. The mean change in earthworm mass (n = 5) from the start of the experiment to the point of destructive sampling (pre-hydration), for earthworms in the constant control (y = 0.735x + 55.836) and drying (y = -1.964x + 77.048) soils. Colours represent each soil type: red = 2.1 (sand), green = 2.2 (sandy loam), blue = 2.4 (loam) and purple = 6S (clay). Error bars show standard error. Dashed line represents no change in mass (y = 0 %).

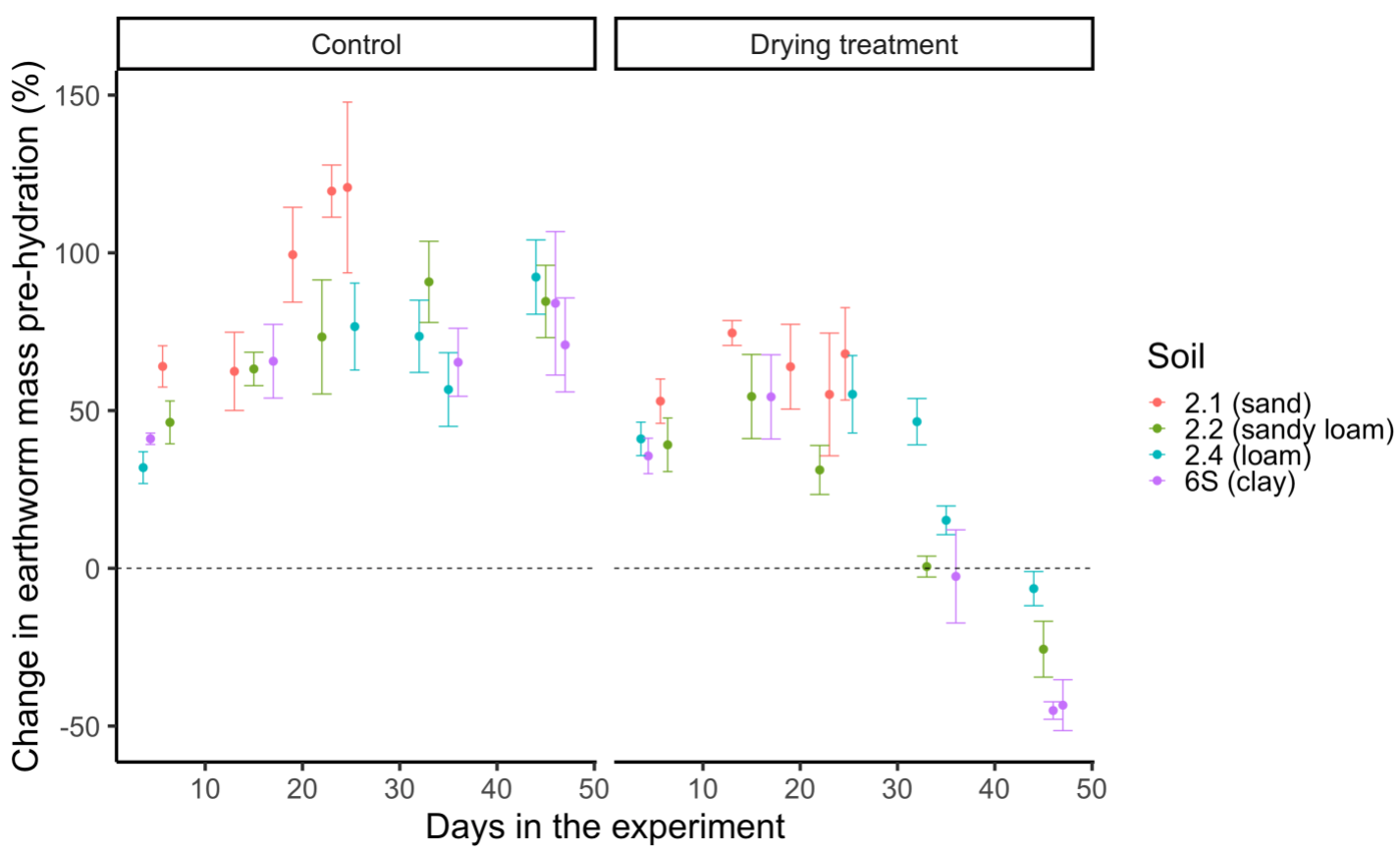

**Figure S9**. The mean change in earthworm mass (n = 5) from the start of the experiment to after 24 hours of hydration, for earthworms in the constant control (y = 0.939x + 16.647) and drying (y = -0.004x + 23.897) soils. Colours represent each soil type: red = 2.1 (sand), green = 2.2 (sandy loam), blue = 2.4 (loam) and purple = 6S (clay). Error bars show standard error.

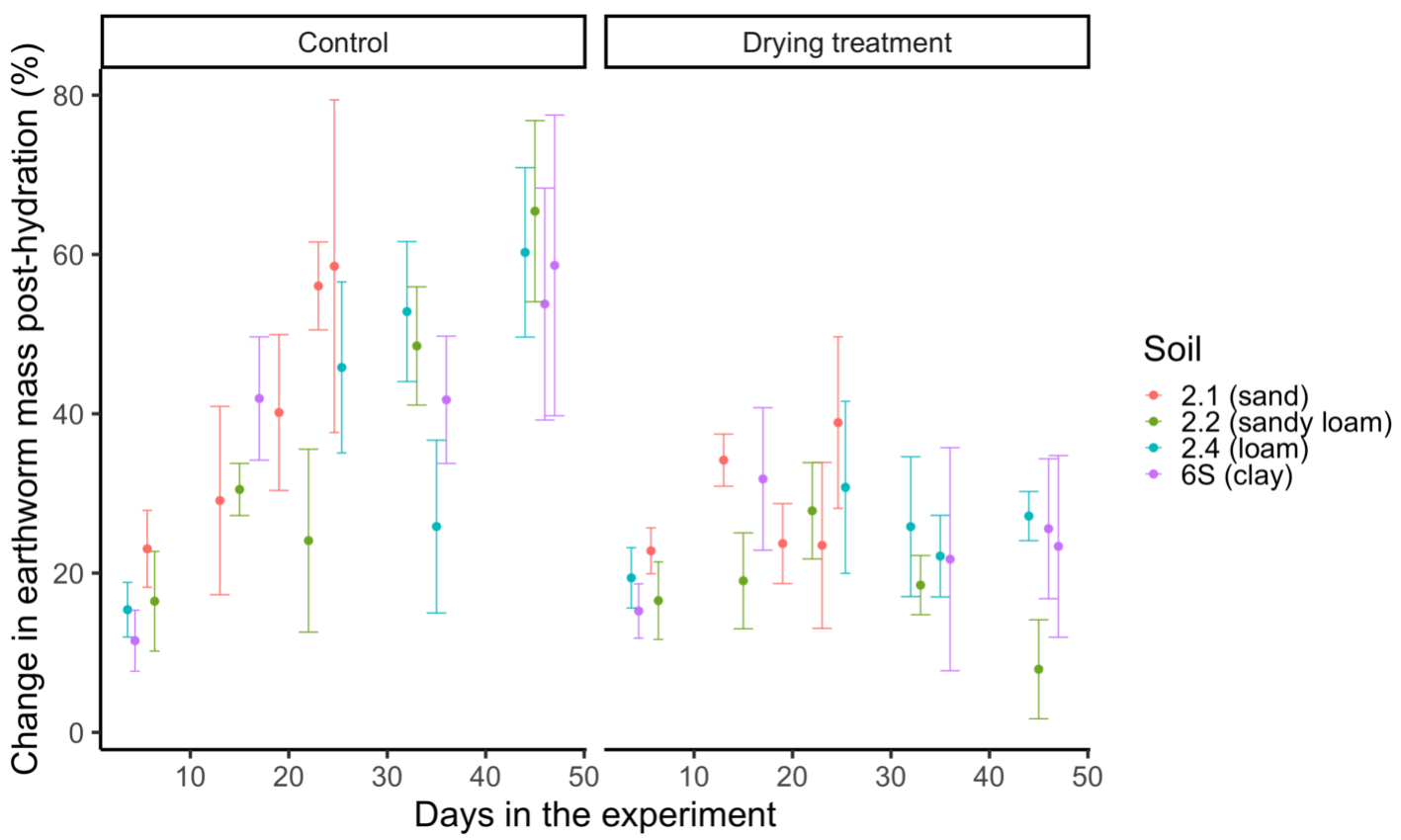

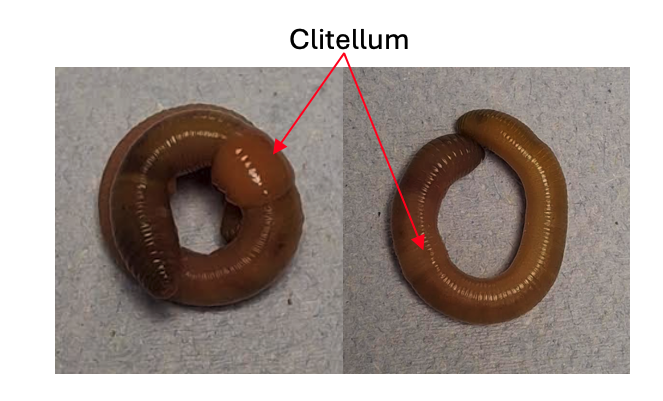

**Figure S10**. Adult *Al. chlorotica* with fully developed (left) and regressed clitellum (right).

**Table S1**. The day of the experiment soils reached each of the sampling points.

| **Soil** | **pF 1.59** | **pF 2.92** | **pF 4.05** | **19.7 wt%** | **15.55 wt%** | **12.39 wt%** |
| --- | --- | --- | --- | --- | --- | --- |
| **2.1** | 6 | 23 | 25 | 6 | 13 | 19 |
| **2.2** | 6 | 33 | 45 | 6 | 15 | 22 |
| **2.4** | 4 | 25 | 44 | 32 | 35 | 44 |
| **6S** | 4 | 17 | 36 | 36 | 46 | 47 |

**Figure S11**. The change in the gravimetric moisture content (%) of each of the four soils over time in the drying conditions. Drying rates were as follows: -0.89 ± 0.008 g day^-1^ for 2.1, -0.87 ± 0.009 g day^-1^ for 2.2, -1.04 ± 0.01 g day^-1^ for 2.4 and -0.95 ± 0.009 g day^-1^ for 6S.

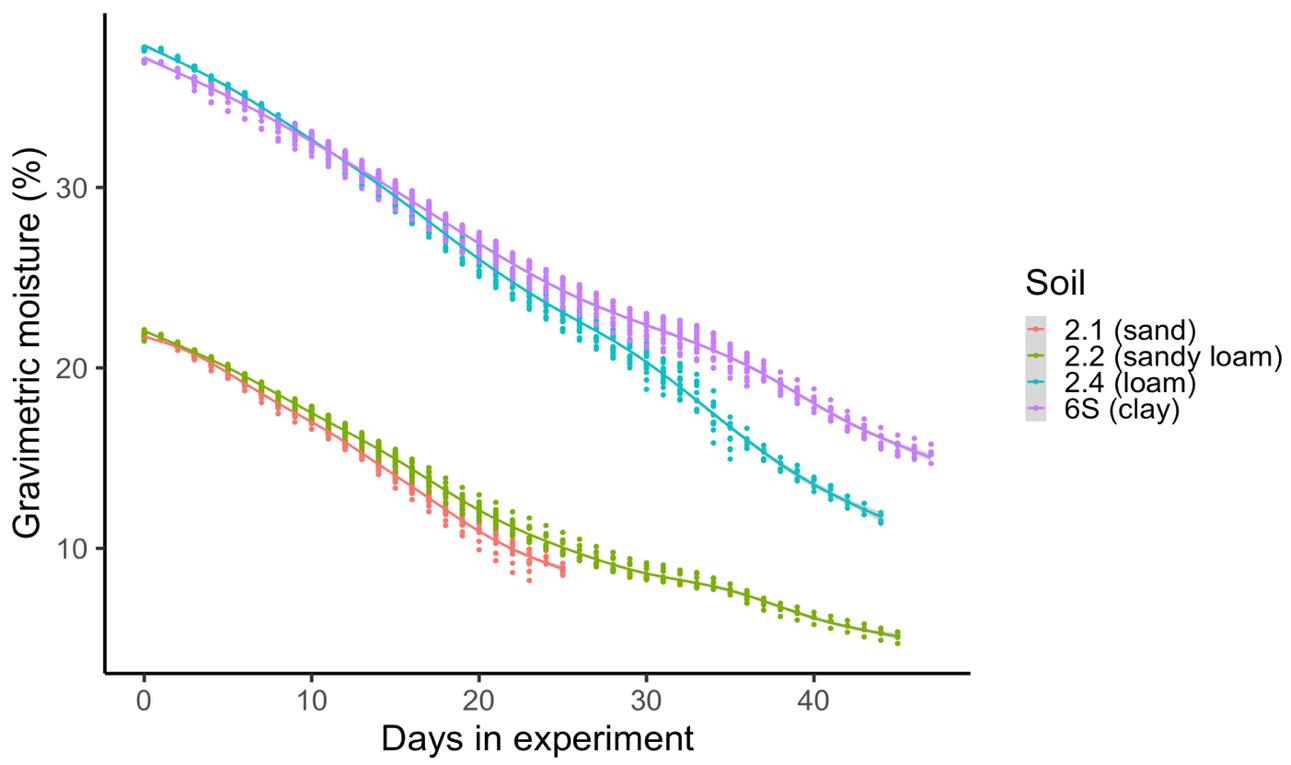
